# Development of a Synthetic Genomics-Based Venezuelan Equine Encephalitis Virus Expressing Secreted *Gaussia* Luciferase for High-Throughput Antiviral Screening

**DOI:** 10.64898/2026.10.21.760855

**Authors:** Miki X Kawano, Akiho Ohkawa, Napatsorn Suwongsaksri, Akatsuki Saito

## Abstract

Venezuelan equine encephalitis virus (VEEV) is a mosquito-borne alphavirus that causes severe neurological disease in humans and animals. Despite its public health importance, no approved antiviral therapy is currently available, and conventional methods for quantifying infectious virus are labor-intensive and poorly suited for high-throughput antiviral screening. Furthermore, research on VEEV is complicated by the restricted availability and international transport of infectious viral materials.

Here, we established a full-length infectious cDNA clone of VEEV entirely from chemically synthesized DNA fragments and rescued recombinant reporter viruses expressing mCherry2, ZsGreen, or the secreted Gaussia luciferase mutant 23 (GLucM23). The reporter viruses retained replication kinetics comparable to those of wild-type VEEV, while the fluorescent reporters enabled direct visualization of infection and virus-induced cytopathic effects. GLucM23-expressing VEEV allowed rapid, sensitive, and non-destructive quantification of viral replication directly from culture supernatants, enabling repeated longitudinal measurements without disrupting infected cells. We further demonstrated the utility of this system by screening approximately 3,000 FDA-approved compounds to identify inhibitors of VEEV replication. Together, these findings establish a versatile synthetic genomics-based reporter virus platform for quantitative studies of VEEV replication and scalable antiviral drug discovery.

## Introduction

Venezuelan equine encephalitis virus (VEEV) is a mosquito-borne alphavirus belonging to the family *Togaviridae* with an 11.5 kb positive-sense single-stranded RNA genome that causes severe neurological disease in both humans and horses [1,2].

VEEV strains are broadly classified into epizootic and enzootic variants based on their genetic lineages, phenotypes, and ecological cycles [3]. While epizootic strains (such as subtypes I-AB and I-C) drive major periodic outbreaks with high mortality in equids and neurological sequelae in humans, enzootic strains (including subtypes I-D, I-E, I-F, and subtypes II–VI) maintain sylvatic persistence across the Americas through continuous circulation between rodent hosts and mosquito vectors. Epizootic outbreaks continue to occur in endemic regions.

In addition to vector-borne transmission, VEEV can also be transmitted through aerosols, and its potential misuse as a biological agent has led to stringent biosafety and biosecurity regulations [4,5]. Despite its medical and veterinary importance, no approved antiviral therapy is currently available, and no vaccine has been licensed for widespread human use. These unmet medical needs highlight the importance of developing efficient experimental platforms for studying VEEV biology and identifying antiviral compounds.

Research on VEEV is challenged not only by the lack of effective therapeutics but also by the limited accessibility of infectious virus. Because VEEV is a highly pathogenic agent, international shipment of infectious viral materials is subject to strict regulatory control, making it difficult for many laboratories to obtain viral isolates. Recent advances in synthetic genomics have enabled the reconstruction of infectious RNA viruses entirely from chemically synthesized DNA, providing an alternative strategy that circumvents the need for international transport of infectious materials. Such reverse genetics approaches also facilitate precise genome engineering and the rapid generation of recombinant viruses for virological research.

Reporter viruses have become indispensable tools for studying virus replication, visualizing infection dynamics, and performing antiviral drug screening. Fluorescent and bioluminescent reporters enable rapid and quantitative assessment of viral replication while supporting high-throughput applications. Among currently available reporter systems, secreted luciferases are particularly attractive because reporter activity can be measured directly from culture supernatants without disrupting infected cells, thereby enabling repeated longitudinal measurements from the same culture. *Gaussia* luciferase mutant 23 (GLucM23), an engineered secreted luciferase with enhanced luminescence intensity and glow-type kinetics [6], is especially well suited for sensitive and scalable antiviral assays. However, no VEEV reporter platform combining synthetic genomics with non-lytic secreted luciferase-based monitoring has been established.

In this study, we established a reverse genetics platform for VEEV by assembling a full-length infectious cDNA clone entirely from chemically synthesized DNA fragments and generated recombinant viruses expressing mCherry2, ZsGreen, or the secreted luciferase GLucM23. We demonstrate that these reporter viruses retain replication characteristics comparable to those of wild-type VEEV while enabling rapid, quantitative, and non-destructive monitoring of viral replication. Furthermore, we applied the GLucM23-expressing VEEV to high-throughput screening of approximately 3,000 FDA-approved compounds, demonstrating the utility of this platform for antiviral drug discovery. Collectively, our study provides a versatile synthetic genomics-based reporter virus platform that facilitates both fundamental studies of VEEV biology and large-scale antiviral screening.

## Materials and Methods

### 2.1. Construction of plasmids expressing fluorescent and luciferase proteins

All experiments involving VEEV infectious cDNA and rescued viruses were performed after obtaining all required institutional approvals. All procedures using infectious viruses were conducted in the biosafety level 3 (BSL-3) laboratory at the University of Miyazaki under appropriate biosafety and biocontainment measures.

A full-length wild-type (WT) VEEV construct was generated based on VEEV strain 3880 (Accession number: L00930.1), which was isolated from a human sample in Panama in 1961, and classified as Subtype I-D (Panamanian genotype) [7]. Because we could not obtain full-length viral RNA of VEEV strain 3880 for biosecurity reasons, seven fragments of the VEEV strain 3880-derived genomic sequence were synthesized by Twist Bioscience (South San Francisco, CA, USA) (**Supplementary Table 1**). We assembled these fragments using PrimeSTAR® GXL DNA Polymerase (TaKaRa, Kusatsu, Japan, Cat# R050A) according to the manufacturer’s instructions. PCR conditions consisted of 35 cycles of 10 s at 98°C, 15 s at 60°C, and 2 min at 68°C, followed by 7 min at 68°C.

The amplified fragment was ligated to a pCMV mammalian expression vector (a kind gift from Dr. Takeshi Kobayashi, The University of Osaka), which encodes the CMV promoter sequence at the 5′ side and the hepatitis delta virus (HDV) ribozyme and the bovine growth hormone (BGH) polyadenylation signal at the 3′ side (**Supplementary Figure 1**) [8], using NEBuilder HiFi DNA Assembly Master Mix (New England Biolabs [NEB], Ipswich, MA, USA, Cat# E2621F) according to the manufacturer’s instructions. The ligated fragments were transformed into NEB 5-alpha F’*I^q^* competent *Escherichia coli* (NEB, Cat# C2992H), and plasmid DNA was purified using the PureYield Plasmid Miniprep System (Promega, Madison, WI, USA, Cat# A1222) according to the manufacturer’s instructions. Multiple clones of each construct were generated and sequence-verified with Sanger sequencing. The sequencing reaction was performed using the SupreDye v3.1 Cycle Sequencing Kit (M&S TechnoSystems, Osaka, Japan, Cat# 063001) and primers (**Supplementary Table 2**) on the Spectrum Compact CE System (Promega).

To generate plasmids encoding reporter-expressing viruses, a 5′ fragment was amplified from the pCMV-VEEV 3880 (WT) plasmid using PrimeSTAR Max DNA Polymerase Ver. 2 (TaKaRa, Cat# R047B) with the primers 3880-1Fver2 (5′- GGTTTAGTGAACCGGATGGGCGGCGCATGAGAGAAGCCCA-3′) and 3880-ZsGreen-R (5′-CACTAGTGACCATTGCTCGCAATTC-3′) according to the manufacturer’s instructions. PCR conditions consisted of 35 cycles of 10 s at 98°C, 5 s at 55°C, and 2 min at 68°C, followed by 7 min at 68°C.

In addition, a 3′ fragment was amplified from the pCMV-VEEV 3880 (WT) plasmid using 3880-ZsGreen-F (5′-CCACCATGTGCCTGCTCGCCAATGT-3′) and 3880-7Rver2 (5′-TGCCATGCCGACCCTGAAATATTAAAAACAAAATCCGATT-3′) primers.

Finally, fragments encoding reporter genes (mCherry2 [9], ZsGreen [10], and GLucM23 [6]) were amplified from synthesized DNA fragments (**Supplementary Table 3**) using 3880-Frag2-F (5′-CAATGGTCACTAGTGACCACCATG-3′) and 3880-Frag2-R (5′-GCAGGCACATGGTGGTCACTAGTGA-3′) primers. These fragments were ligated using NEBuilder HiFi DNA Assembly Master Mix. The plasmids were amplified as described above.

### 2.2. Cell culture

Human hepatocyte-derived Huh7 cells (Japanese Collection of Research Bioresources Cell Bank [JCRB], Ibaraki, Japan, Cat# JCRB0403) were maintained in Dulbecco’s modified Eagle medium (DMEM; Nacalai Tesque, Kyoto, Japan, Cat# 08458-16) supplemented with 10% fetal bovine serum (FBS; Cytiva, Marlborough, MA, USA, Cat# SH30396) and 1× penicillin-streptomycin (Pe/St; Nacalai Tesque, Cat# 09367-34) at 37°C in a humidified incubator with 5% CO_2_.

African green monkey kidney-derived Vero cells (JCRB, Cat# JCRB0111) were maintained in Minimum Essential Medium (MEM) with Earle’s salts and L-glutamine (Nacalai Tesque, Cat# 21442-25) supplemented with 10% FBS, Pe/St, 1× MEM non-essential amino acids solution (Nacalai Tesque, Cat# 06344-56), and 20 mM HEPES buffer (Nacalai Tesque, Cat# 17557-94) at 37°C in a humidified incubator with 5% CO_2_.

Human neuroblastoma-derived NB-1 cells (JCRB, Cat# IFO50295) were maintained in RPMI 1640 medium supplemented with L-glutamine (Nacalai Tesque, Cat# 30264-56), 10% FBS, and Pe/St at 37°C in a humidified incubator with 5% CO_2_.

### 2.3. Rescue and propagation of virus stocks

Huh7 cells were plated in a 12-well plate (Wako, Cat# 636-28421) at 8.3 × 10^5^ cells/well and incubated overnight. Cells were transfected with plasmids using Lipofectamine 3000 transfection reagent (Thermo Fisher Scientific, Waltham, MA, USA, Cat# L3000015) in Opti-MEM I Reduced Serum Medium (Thermo Fisher Scientific, Cat# 31985062). Culture supernatants were harvested at 5 days post-transfection.

For virus propagation, Vero cells were seeded in a T75 flask at 3 × 10^6^ cells per flask and incubated overnight. VEEV 3880-mCherry2, VEEV 3880-ZsGreen, VEEV 3880-GLucM23, and WT VEEV 3880 were diluted in medium, and cells were infected at an MOI of 0.001. After adsorption for 2 h at 37°C, the inoculum was removed, and 20 mL of fresh MEM supplemented with 2% FBS and Pe/St was added. Infected cultures were incubated at 37°C for 5 days.

At 5 days post-infection, culture supernatants were collected and transferred to 15 mL tubes. The supernatants were clarified by centrifugation at 3,000 rpm for 10 min at room temperature to remove cellular debris, then transferred to new 15 mL tubes. To remove cell-free GLucM23 from the virus-containing supernatants, one volume of 40% polyethylene glycol #6,000 (Nacalai Tesque, Cat# 28254-85) solution was mixed with four volumes of clarified supernatant, then incubated overnight at 4°C. The samples were then centrifuged at 3,000 rpm for 40 min at 4°C. After carefully removing the supernatant, the resulting pellets were resuspended in DMEM supplemented with 10% FBS and Pe/St. The prepared virus-containing samples were aliquoted into cryovials and stored at −80°C until use.

### 2.4. Plaque assay

Vero cells were plated in a 24-well plate (Wako, Cat# 630-28441) at 8 × 10^4^ cells/well and incubated overnight. Virus stocks were serially 10-fold diluted in serum-free MEM, and 250 µL of each dilution was added to the cell monolayers. After adsorption for 1 h at 37°C, the cells were overlaid with 500 µL of MEM (Thermo Fisher Scientific, Cat# 11935046) containing 3% FBS, Pe/St, and 1.5% carboxymethylcellulose sodium salt (Sigma-Aldrich, Cat# C4888-500G).

At 4 days post-infection, cells were fixed and stained with crystal violet. Plaques were counted, and virus titers were calculated as plaque-forming units (PFU)/mL. The VEEV constructs analyzed included VEEV 3880-mCherry2, VEEV 3880-ZsGreen, VEEV 3880-GLucM23, and WT VEEV 3880.

### 2.5. Measurement of GLucM23 activity

Vero cells were plated in a poly-D-lysine-coated 96-well plate (Corning, Cat# 356461) at 2 × 10^4^ cells/well and incubated overnight. For infection, VEEV 3880-GLucM23 and WT VEEV 3880 were diluted in infection medium to obtain the indicated multiplicities of infection (MOIs) (MOIs of 0.005 or 0.001). Virus inoculum was added to the cells, and plates were incubated at 37°C for 1 h to allow virus adsorption. Cells were then washed twice with fresh infection medium and maintained in 150 µL of fresh medium. Culture supernatants were collected at 0, 24, and 48 h post-infection (hpi).

GLucM23 activity in culture supernatants was measured using the Pierce *Gaussia* Luciferase Glow Assay Kit (Thermo Fisher Scientific, Cat# 16160) according to the manufacturer’s instructions. *Gaussia* Glow Assay Buffer was mixed with coelenterazine substrate at a 1:100 dilution before use. Five microliters of each sample were mixed with 5 µL of 2× Cell Lysis Buffer in a white 96-well plate, followed by the addition of 25 µL of substrate solution. Relative light units (RLU) were measured immediately using a luminometer.

### 2.6. Quantification of viral RNA by qRT-PCR

Vero cells were plated in a poly-D-lysine-coated 96-well plate at 2 × 10^4^ cells/well and incubated overnight. Cells were infected with VEEV 3880-mCherry2, VEEV 3880-ZsGreen, VEEV 3880-GLucM23, or WT VEEV 3880 at an MOI of 0.001 or 0.005, as described above. Culture supernatants were collected at 0, 24, and 48 hpi for viral RNA quantification.

The culture supernatant was mixed with 2× RNA lysis buffer (2% Triton X-100, 50 mM KCl, 100 mM Tris-HCl (pH 7.4), 40% glycerol, and 0.4 U/µL Recombinant RNase Inhibitor (TaKaRa, Cat# 2313A)) [11]. qRT-PCR was performed using a One Step TB Green PrimeScript PLUS RT-PCR Kit (Perfect Real Time) (TaKaRa, Cat# RR096A) with primers 3880-qRT-PCR-F (5′-TTTCTCTGCACCCACAGTTC-3′) and 3880-qRT-PCR-R (5′- CTGACGGACCACTTCCTTATTC-3′) according to the manufacturer’s instructions. The PCR protocol was 42°C for 5 min, 95°C for 10 s, 40 cycles of 95°C for 5 s, and 60°C for 34 s. Viral RNA copy numbers were calculated from standard curves generated using known concentrations of the pCMV-VEEV 3880 plasmid.

### 2.7. Western blot analysis

Vero cells were plated in a poly-D-lysine-coated 24-well plate (Corning, Cat# 356414) at a density of 8 × 10^4^ cells/well and incubated overnight. Cells were infected with VEEV 3880-mCherry2, VEEV 3880-ZsGreen, VEEV 3880-GLucM23, or WT VEEV 3880 at MOIs of 0.001 or 0.005. After virus adsorption for 1 h at 37°C, cells were washed twice with 500 µL of fresh medium and maintained in 500 µL of fresh infection medium. At 24 hpi, cells were harvested into 1.5 mL tubes and centrifuged at 6,000 rpm for 1 min. After removal of the supernatant, cell pellets were washed with PBS twice and lysed in 100 µL of 2× Bolt LDS Sample Buffer (Thermo Fisher Scientific, Cat# B0008) containing 2% β-mercaptoethanol (Bio-Rad, Hercules, CA, USA, Cat# 1610710). Samples were incubated at 70°C for 10 min to denature proteins.

Protein expression was analyzed using a Simple Western Abby system (ProteinSimple, San Jose, CA, USA) with a 12–230 kDa Separation Module, 8 × 25 capillary cartridge (ProteinSimple, Cat# SM-W004A) according to the manufacturer’s instructions. VEEV non-structural protein 1 (nsP1) was detected using an anti-VEEV nsP1 rabbit monoclonal antibody (HL1472; GeneTex, Cat# GTX636946). mCherry expression was detected using an anti-mCherry rabbit monoclonal antibody (E5D8F; Cell Signaling Technology, Cat# 43590S). The amount of input protein was measured using the Total Protein Detection Module (ProteinSimple, Cat# DM-TP01).

### 2.8. Antiviral inhibitor assay

For the antiviral assay in Vero cells, cells were treated with serially diluted recombinant human IFN-β (PeproTech, Cat# 300-02BC) for 24 h prior to infection. Cells were infected with VEEV 3880-GLucM23 virus (MOI 0.001). After 1 h of virus adsorption at 37°C, the inoculum was removed, and cells were cultured in fresh medium. To measure antiviral activity, culture supernatants were collected at 24 and 48 hpi and analyzed as described above. Luminescence values were used to evaluate the inhibitory activity against VEEV compared with the untreated negative control.

For the primary screening in NB-1 cells, cells were plated in a poly-D-lysine-coated 96-well plate at 2 × 10⁴ cells/well and incubated overnight. Cells were infected with VEEV 3880-GLucM23 virus (MOI 0.001). After 1 h of virus adsorption at 37°C, the inoculum was removed, and cells were cultured in medium containing test compounds or controls. The inhibitor screen used an FDA-approved Drug Library (Selleck, Cat# L1300) containing approximately 3,000 drugs (n = 1). β-D-N⁴-hydroxycytidine (Cell Signaling Technology, Cat# 81178) and dimethyl sulfoxide (DMSO) were used as positive and negative controls, respectively. The antiviral activity was evaluated at 48 hpi as described above.

For the secondary screening, drugs that reduced luminescence to less than 20% of the DMSO control were selected. NB-1 cells were infected (MOI 0.001) and treated with serially diluted drugs as described above (n = 4).

### 2.9. Cytotoxicity assay of candidate drugs

NB-1 cells were plated in a poly-D-lysine-coated 96-well plate at 2 × 10⁴ cells/well and incubated overnight. Cells were cultured in medium containing serially diluted test compounds or controls (n = 4). After 48 h, the cytotoxicity of the test compounds was evaluated with the Cell Counting Kit-8 (Dojindo, Cat# CK04) according to the manufacturer’s instructions.

### 2.10. Data and statistical analysis

Data are presented as the mean ± SD of technical replicates from one assay and are representative of at least three independent experiments. The 50% effective concentration (EC₅₀) and 50% cytotoxic concentration (CC₅₀) values were calculated by nonlinear regression analysis. All data analyses were performed using GraphPad Prism version 11.0.0 (GraphPad Software, Boston, MA, USA).

## Results

### 3.1. Construction and rescue of VEEVs expressing reporter proteins

VEEV reporter constructs were generated by multi-fragment DNA assembly using the pCMV-VEEV 3880 backbone. Based on a previous report [12], the reporter genes encoding mCherry2, ZsGreen, or GLucM23 were inserted into the region between the C and PE2 genes via a T2A peptide sequence (**Supplementary Figure 1**). Reporter viruses were rescued by transfecting Huh7 cells with the plasmids.

Clear cytopathic effects (CPE) were observed in Huh7 cells transfected with each VEEV construct, including the WT and reporter virus constructs (**Supplementary Figure 2**). Robust fluorescence was detected in cells transfected with the mCherry2- or ZsGreen-expressing constructs, which exhibited strong red and green fluorescence, respectively. In contrast, cells transfected with WT VEEV or the GLucM23-expressing construct showed typical CPE but no detectable fluorescence (**Supplementary Figure 2**). We performed a plaque assay in Vero cells to determine the viral titers of the reporter viruses. The infectious titers ranged from 1.8 × 10⁵ to 3.8 × 10⁵ PFU/mL. These findings indicate successful rescue of infectious VEEV from the plasmid constructs and confirm reporter protein expression in the fluorescent reporter viruses.

### 3.2. Reporter gene insertion has minimal effects on viral replication

To evaluate the impact of reporter gene insertion on viral replication capacity, we measured viral RNA levels in the culture supernatants of infected Vero cells. The results demonstrated that VEEV expressing reporter proteins replicated efficiently in Vero cells at MOIs of both 0.005 and 0.001, with replication efficiency comparable to that of the WT virus (**Figure 1A**), suggesting that the insertion of reporter proteins minimally affected viral replication. Fluorescence microscopy observations confirmed robust expression of mCherry2 and ZsGreen in infected Vero cells (**Figure 1B**). Furthermore, observations in human neuroblastoma-derived NB-1 cells at 15 and 24 hpi demonstrated that both VEEV 3880-mCherry2 and VEEV 3880-ZsGreen showed positive fluorescence signals at 15 hpi (**Supplementary Figure 3A**), with nearly all infected cells becoming positive at 24 hpi (**Supplementary Figure 3B**).

**Figure 1.**
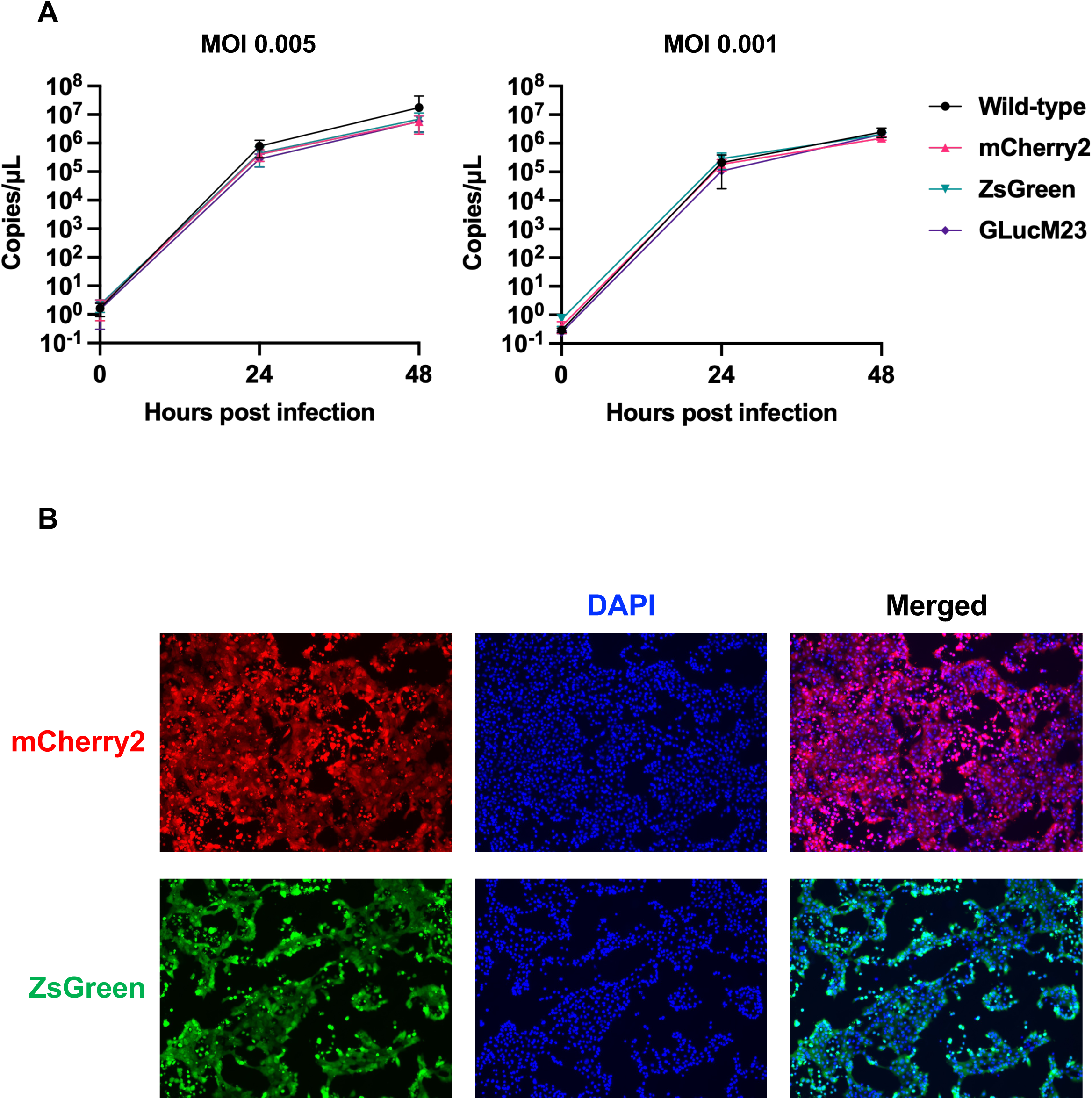
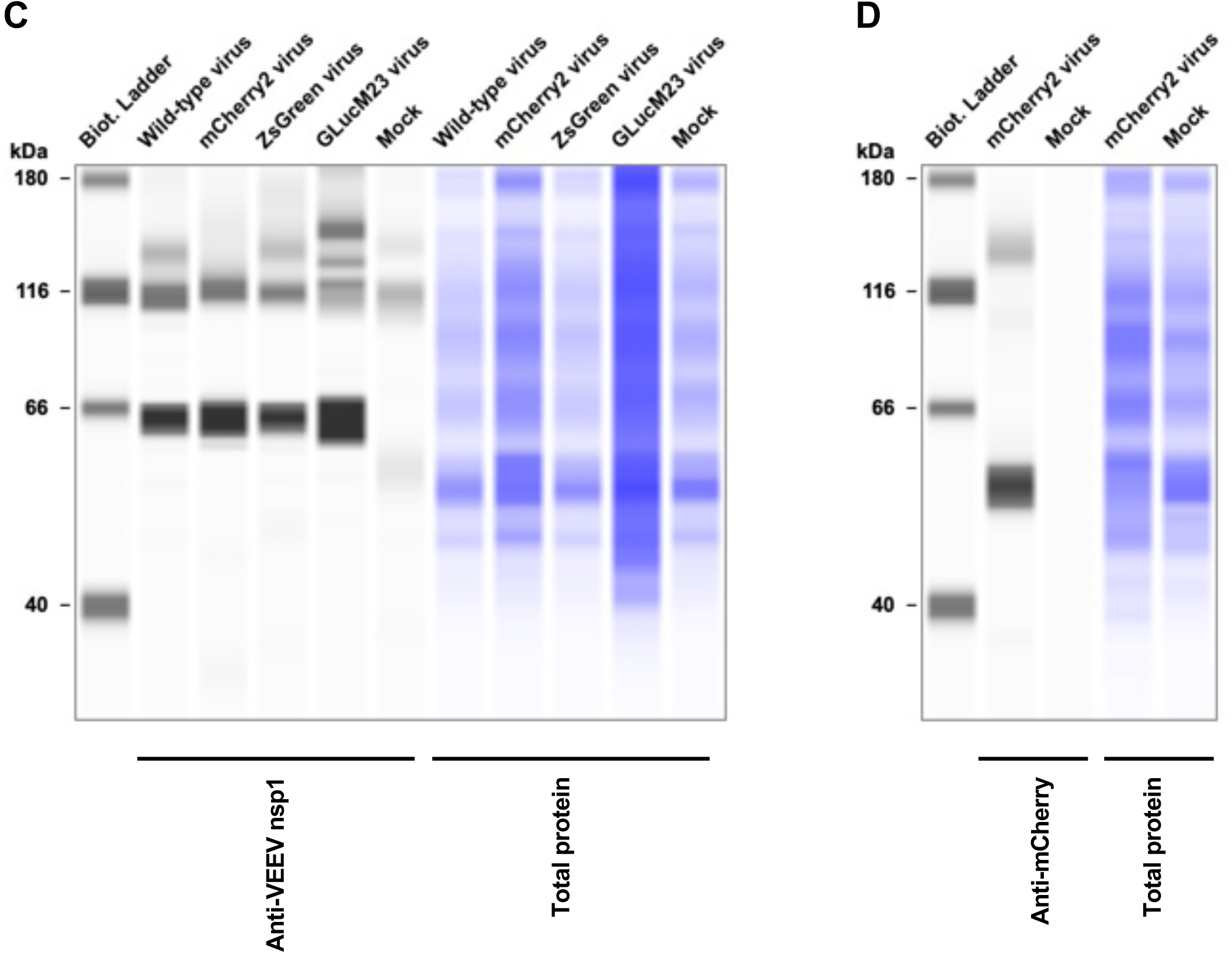
Reporter gene insertion minimally affects VEEV replication and reporter protein expression. (**A**) Replication kinetics of WT VEEV and reporter-expressing VEEV (mCherry2, ZsGreen, and GLucM23) in Vero cells infected at MOIs of 0.005 and 0.001. Viral RNA in culture supernatants was quantified by qRT-PCR at the indicated time points. Data are presented as the mean ± SD. (**B**) Fluorescence microscopy of Vero cells infected with VEEV 3880-mCherry2 or VEEV 3880-ZsGreen. Cells were counterstained with DAPI (blue). Representative fluorescence, DAPI, and merged images are shown. (**C**) Western blot analysis of VEEV nsP1 protein expression in Vero cells harvested at 24 h post-infection (hpi). Total protein staining was used as the loading control. (**D**) Western blot analysis of mCherry protein expression in Vero cells harvested at 24 hpi. Total protein staining was used as the loading control.

Western blot analysis showed that all reporter viruses expressed viral nsP1 at levels comparable to those observed in cells infected with WT VEEV (**Figure 1C**). In addition, mCherry protein was specifically detected in cells infected with VEEV 3880-mCherry2 (**Figure 1D**), confirming functional expression of the inserted reporter gene. These findings suggest that reporter gene insertion did not markedly compromise viral protein synthesis and support the use of these engineered VEEV strains as tools for virological studies and antiviral screening.

### 3.3. Real-time monitoring of VEEV replication using secreted luciferase

We next evaluated whether the GLucM23-expressing VEEV could be used for rapid and non-lytic quantification of viral replication. Because GLucM23 is secreted into the culture supernatant, luciferase activity can be measured without lysing infected cells, allowing repeated monitoring of viral replication over time. Vero cells were infected with VEEV 3880-GLucM23 at two different multiplicities of infection (MOIs of 0.005 or 0.001), and culture supernatants were collected at 0, 24, and 48 hpi for luciferase activity measurement (**Figures 2A and 2B**).

**Figure 2.**
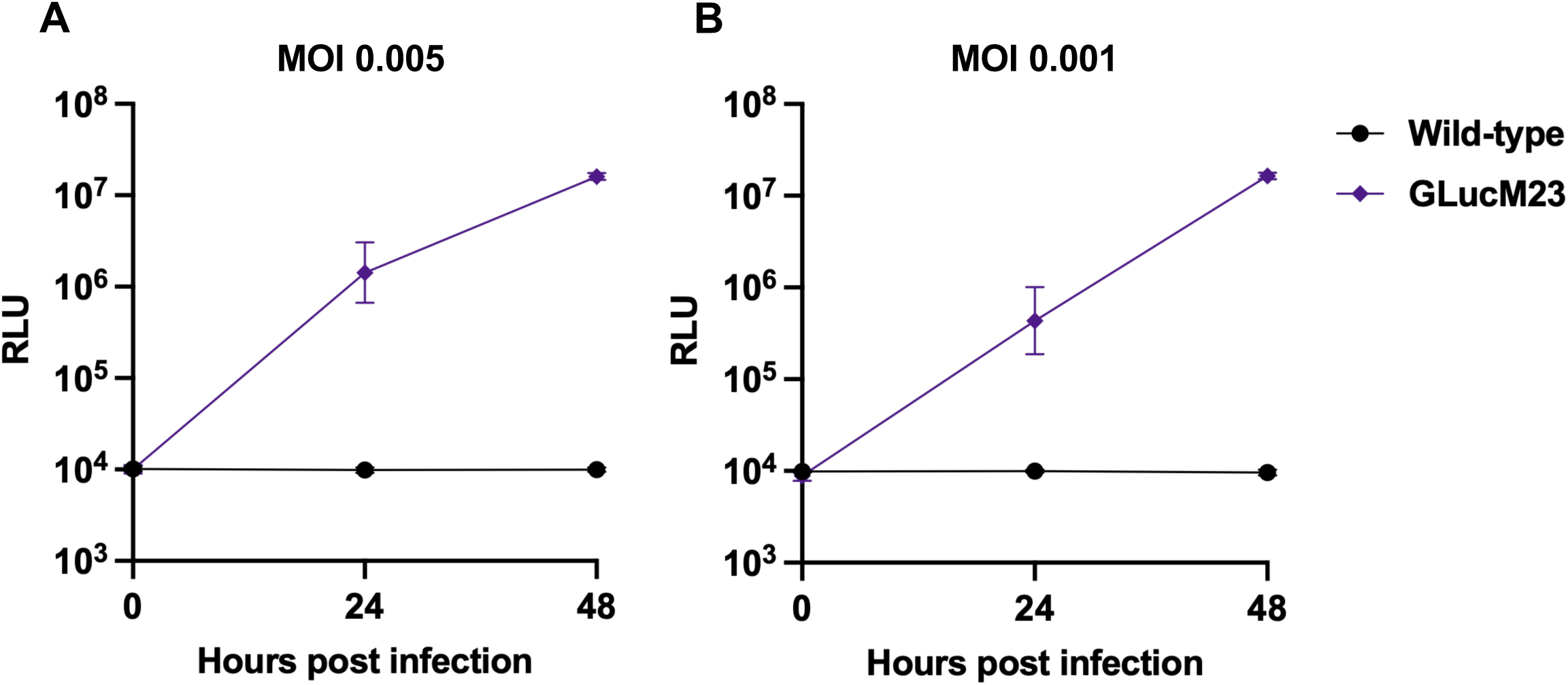
Real-time monitoring of VEEV replication using secreted GLucM23. Vero cells were infected with VEEV 3880-GLucM23 or WT VEEV at MOIs of 0.005 (**A**) and 0.001 (**B**). Culture supernatants were collected at the indicated time points, and *Gaussia* luciferase activity was measured directly from the culture supernatants without cell lysis. The results, presented as the mean ± SD of sextuplicate measurements from one assay, are representative of at least three independent experiments.

Luciferase activity increased by more than 1,000-fold within 48 hpi at both MOIs. In contrast, the WT virus-infected control showed only minimal background activity, confirming the specificity of the GLucM23 signal. These results demonstrate that secreted luciferase activity provides a sensitive and convenient readout for monitoring VEEV replication kinetics.

### 3.4. Application of GLucM23-expressing VEEV to high-throughput antiviral screening

To assess the applicability of the GLucM23-expressing VEEV system for antiviral assays, we first treated Vero cells with serially diluted recombinant human IFN-β prior to infection with VEEV 3880-GLucM23. IFN-β treatment inhibited GLucM23 activity in the culture supernatant in a dose-dependent manner at both 24 and 48 hpi (**Supplementary Figures 4A and 4B**), yielding a half-maximal effective concentration (EC₅₀) of 0.0104 U/mL at 48 hpi (**Figure 3A**). Next, to evaluate the utility of the GLucM23-expressing VEEV system for high-throughput antiviral screening, we tested approximately 3,000 FDA-approved compounds using NB-1 cells derived from human neuroblastoma. Antiviral activity was evaluated by measuring the reduction in GLucM23 activity in the culture supernatant at 48 hpi.

**Figure 3.**
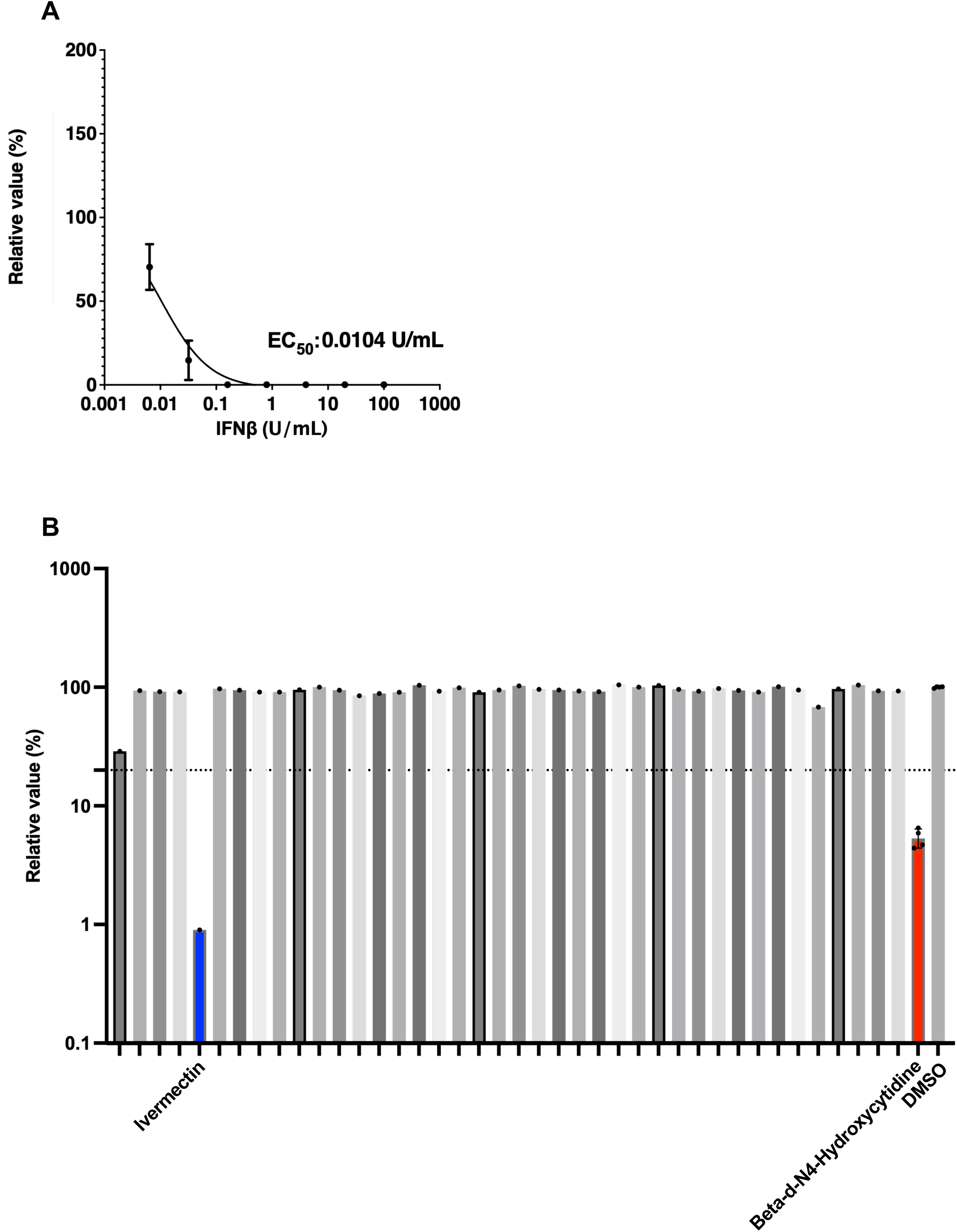
Application of the VEEV 3880-GLucM23 reporter virus to antiviral assays. (**A**) Vero cells were treated with serially diluted recombinant human IFN-β for 24 h prior to infection. Cells were infected with VEEV 3880-GLucM23 (MOI 0.001), and the antiviral activity was determined at 48 hpi by measuring *Gaussia* luciferase activity in the culture supernatant. Relative values (%) normalized to the untreated control and the calculated EC₅₀ value are shown. The results, presented as the mean ± SD of sextuplicate measurements from one assay, are representative of at least three independent experiments. (**B**) Representative results from the primary screen in NB-1 cells. NB-1 cells were infected with VEEV 3880-GLucM23 virus (MOI 0.001). After 1 h of virus adsorption at 37°C, the cells were washed and cultured in medium containing test compounds or controls. β-D-N⁴-hydroxycytidine (highlighted in red) and dimethyl sulfoxide (DMSO) were used as positive and negative controls, respectively. The antiviral activity was determined at 48 hpi by measuring *Gaussia* luciferase activity in the culture supernatant. Ivermectin is highlighted in blue. The dashed line indicates the hit cutoff (20% of the DMSO-treated control).

β-D-N⁴-hydroxycytidine, the positive control in this assay, showed a marked reduction in luciferase activity compared with the DMSO-treated negative control (**Figure 3B**), validating the assay performance. Among the tested compounds, several reduced the relative GLucM23 signal below the predefined cutoff value of 20% compared with the DMSO-treated negative control (**Figure 3B**).

Among approximately 3,000 drugs, 31 drugs suppressed VEEV replication to less than 20% of the DMSO control (**Table 1**). Furthermore, ivermectin showed a strong reduction in reporter activity (**Figure 3B and Table 1**), consistent with a previous study that showed ivermectin’s ability to suppress VEEV replication [13]. In addition, the candidate list included several proteasome inhibitors (bortezomib, carfilzomib, and ixazomib) and the selective inhibitor of nuclear export, selinexor (KPT-330) (**Table 1**), which target host pathways previously shown to be critical for VEEV replication [14,15]. However, when the 31 candidate drugs identified in the primary screen (such as carfilzomib and auranofin) were evaluated in a secondary screen using serial dilutions, their EC₅₀ values (0.0356 µM and 0.3085 µM, respectively) were close to their 50% cytotoxic concentration (CC₅₀) values (0.09657 µM and 1.103 µM, respectively), indicating that no novel compounds with selective anti-VEEV activity and low cytotoxicity were identified from this library (**Supplementary Figure 5**).

**Table 1.** List of candidate drugs. Drugs that suppressed VEEV replication to less than 20% of the DMSO control were used for the secondary screening.

| Drug | Selleck Cat# | Relative value (%) |
| --- | --- | --- |
| Bortezomib (PS-341) | S1013 | 6.5 |
| Paclitaxel (NSC 125973) | S1150 | 13.1 |
| Vincristine (NSC-67574) sulfate | S1241 | 15.5 |
| Ivermectin (MK-933) | S1351 | 0.9 |
| Mycophenolate mofetil (RS-61443) | S1501 | 11.5 |
| Crystal violet | S1917 | 18.8 |
| Ixazomib (MLN2238) | S2180 | 13.5 |
| Ixazomib citrate (MLN9708) analogue | S2181 | 13.0 |
| (S)-10-Hydroxycamptothecin | S2423 | 10.2 |
| Mitoxantrone (NSC-301739) 2HCl | S2485 | 13.7 |
| Carfilzomib (PR-171) | S2853 | 3.6 |
| Hydroxycamptothecin | S3898 | 8.2 |
| Auranofin | S4307 | 7.3 |
| Doramectin | S5117 | 8.1 |
| Saikosaponin D | S5454 | 1.9 |
| Lanatoside C | S5554 | 0.6 |
| Pyrvinium | S5816 | 0.9 |
| Selinexor (KPT-330) | S7252 | 3.2 |
| Puromycin (CL13900) 2HCl | S7417 | 0.2 |
| Monomethyl auristatin E (MMAE) | S7721 | 19.5 |
| Combretastatin A4 | S7783 | 14.2 |
| Docetaxel trihydrate | S7787 | 17.9 |
| Harringtonine | S9063 | 0.1 |
| Belotecan (CKD-602) hydrochloride | S6631 | 13.5 |
| Homoharringtonine (CGX-635) | S9015 | 0.1 |
| Staurosporine (AM-2282) | S1421 | 0.2 |
| Ixazomib citrate (MLN9708) | S4432 | 14.5 |
| Deslanoside | E0680 | 0.2 |
| Gramicidin | E2970 | 0.1 |
| Melittin | S9794 | 4.1 |
| Camptothecin (NSC-100880) | S1288 | 15.5 |

## Discussion

In this study, we developed and characterized synthetic genomics-based VEEV reporter viruses expressing either a secreted luciferase (GLucM23) or fluorescent proteins (mCherry2 or ZsGreen) to facilitate the monitoring of viral replication. Our findings clearly demonstrate that the introduction of these reporter proteins does not have a detrimental effect on viral replication kinetics or viral protein expression. Both secreted luciferase activity and viral RNA levels increased in parallel, confirming that these reporter constructs accurately reflect viral replication dynamics.

The use of these reporter viruses offers significant advantages for virological studies. First, the secreted GLucM23 system allows for rapid, sensitive, and non-lytic monitoring of viral replication directly from culture supernatants (**Figure 2**), while the fluorescent reporter viruses enable direct visualization of infected cells and virus-induced cytopathic effects by fluorescence microscopy (**Figure 1B**). The strong correlation between the reporter signal and viral RNA copy numbers validates their reliability as surrogate markers for viral load (**Figures 1A and 2**). Second, the GLucM23-expressing VEEV provides a time-efficient and scalable platform for high-throughput antiviral screening. After confirming the sensitivity of this system through the dose-dependent inhibition of viral replication by recombinant human IFN-β in Vero cells (**Figure 3A**), we screened an FDA-approved drug library of approximately 3,000 compounds in human neuroblastoma-derived NB-1 cells to evaluate its applicability to drug discovery (**Figure 3B**). Consistent with a previous report [13], ivermectin demonstrated significant anti-VEEV activity alongside the positive control β-D-N⁴-hydroxycytidine, validating the reliability of our assay system. Moreover, our primary screen successfully identified proteasome inhibitors, including bortezomib, carfilzomib, and ixazomib, as well as the selective nuclear export inhibitor selinexor (KPT-330) (**Table 1**). Because previous studies have demonstrated that bortezomib-mediated inhibition of the ubiquitin–proteasome system and selective inhibitors of nuclear export (SINE) targeting CRM1/XPO1 potently suppress VEEV replication [14,15], the identification of these mechanistically relevant compounds further corroborates the robustness of our screening platform.

However, this study has several limitations. First, although 31 initial hit compounds were identified in the primary screen, several had previously been reported, and secondary dose– response evaluations revealed that the EC₅₀ values of the candidates were close to their CC₅₀ values; thus, we did not identify novel compounds exhibiting both potent anti-VEEV activity and low cytotoxicity from the library used in this study (**Supplementary Figure 5**). Nevertheless, our results clearly demonstrate the utility of this platform for high-throughput screening, and future studies screening larger and more diverse chemical libraries may lead to the discovery of novel anti-VEEV therapeutics. Second, because the reporter-expressing VEEV developed in this study is based on a single enzootic strain (strain 3880, subtype I-D), future studies should extend this reverse genetics platform to other VEEV subtypes, including epizootic strains.

Beyond antiviral drug screening, the GLucM23-expressing VEEV reporter virus system has potential applications in vaccine development and immunological studies. The rapid, quantitative, and non-lytic nature of this platform makes it well-suited for evaluating vaccine-induced immune responses. For instance, this system could be applied to measure neutralizing antibody responses elicited by different VEEV vaccine candidates, enabling rapid assessment and comparison of vaccine efficacy without relying on labor-intensive plaque or TCID₅₀ assays. Furthermore, the flexibility of our synthetic genomics approach circumvents the need for international transport of infectious viral materials and facilitates the generation of recombinant vaccine vectors expressing heterologous antigens or attenuated derivatives suitable for vaccine development.

In summary, our study demonstrates that reporter viruses expressing secreted luciferase or fluorescent proteins are reliable and effective tools for monitoring VEEV replication dynamics. These reporter constructs minimally impact viral replication and enable rapid, high-throughput screening of antiviral compounds. The ability to easily track viral infection and cytopathic effects over time greatly enhances both basic research and drug discovery efforts. Overall, these reporter viruses represent a valuable platform for advancing our understanding of VEEV replication and accelerating the development of novel antiviral therapies and vaccines.

## Acknowledgements

The authors thank Dr. Takeshi Kobayashi (The University of Osaka) for providing the pCMV vector. The authors thank Ms. Tomoko Nishiuchi, Ms. Natsumi Matsubara, and the staff of CADIC, University of Miyazaki, for their assistance.

## Disclosing the Use of Artificial Intelligence (AI)

During the preparation of this manuscript, the authors used Gemini (1.131.4.1198) solely for English language editing and proofreading of the text originally drafted by the authors. No AI tools were used for data analysis, interpretation, or the generation of scientific content. After using this tool, the authors carefully reviewed and edited the content as needed and take full responsibility for the accuracy, integrity, and originality of the published work.

## Supporting information

**Supplementary Figure 1. Schematic representation of reporter-expressing VEEV constructs.**

Schematic diagrams of the recombinant VEEV genomes used in this study. Reporter genes encoding mCherry2, ZsGreen, or GLucM23 were inserted between the capsid (C) and PE2 coding regions via a Thosea asigna virus 2A (T2A) peptide sequence. The full-length infectious cDNA clone was assembled in a pCMV vector containing a CMV promoter at the 5′ end and an HDV ribozyme followed by a BGH polyadenylation signal at the 3′ end.

**Supplementary Figure 2. Rescue of reporter-expressing VEEV from infectious cDNA clones.**

Representative bright-field and fluorescence images of mock-transfected Huh7 cells or Huh7 cells at 5 days post-transfection with plasmids encoding WT VEEV or reporter-expressing VEEV (mCherry2, ZsGreen, or GLucM23).

**Supplementary Figure 3. Visualization of reporter protein expression in VEEV-infected NB-1 cells.**

(**A**) Representative bright-field and fluorescence images of mock-infected NB-1 cells or NB-1 cells infected with WT VEEV, VEEV 3880-mCherry2, or VEEV 3880-ZsGreen at 15 h post-infection (hpi).

(**B**) Representative bright-field and fluorescence images of mock-infected NB-1 cells or NB-1 cells infected with the indicated viruses at 24 hpi.

**Supplementary Figure 4. Evaluation of anti-VEEV activity of human IFN-β using VEEV 3880-GLucM23.**

Vero cells were treated with serially diluted recombinant human IFN-β for 24 h prior to infection. Cells were infected with VEEV 3880-GLucM23 (MOI 0.001), and the antiviral activity was determined at 24 hpi (**A**) and 48 hpi (**B**) by measuring *Gaussia* luciferase activity in the culture supernatant. The results, presented as the mean ± SD of sextuplicate measurements from one assay, are representative of at least three independent experiments.

**Supplementary Figure 5. Representative results of the secondary screening.**

NB-1 cells were infected with VEEV 3880-GLucM23 virus (MOI 0.001) and treated with serially diluted candidate compounds (carfilzomib and auranofin). After 48 h, cell viability (left panels, CC₅₀) and antiviral activity (right panels, EC₅₀) were evaluated and expressed as relative values (%) compared with the DMSO-treated control. Data are presented as the mean ± SD of quadruplicate measurements.

