## Supplemental Materials for "Development of a Synthetic Genomics-Based Venezuelan Equine Encephalitis Virus Expressing Secreted *Gaussia* Luciferase for High-Throughput Antiviral Screening"

**Supplementary Figure 1**

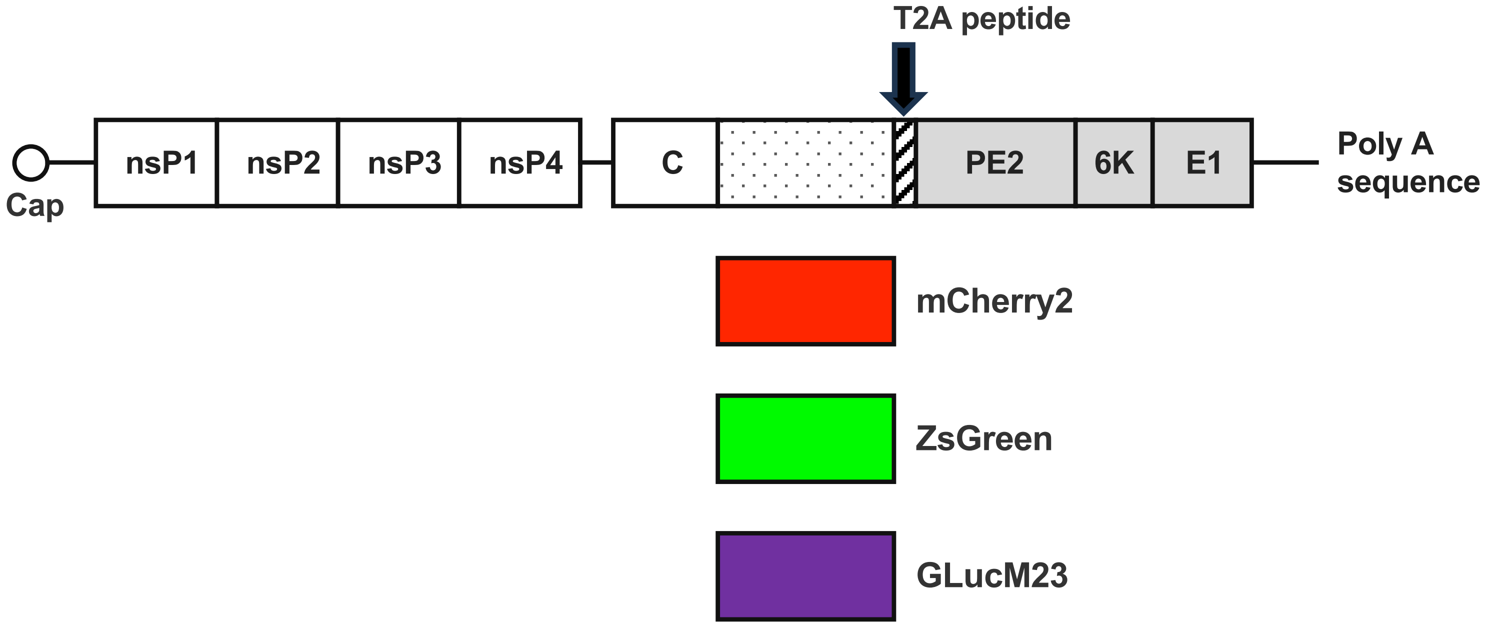

### Supplementary Figure 2

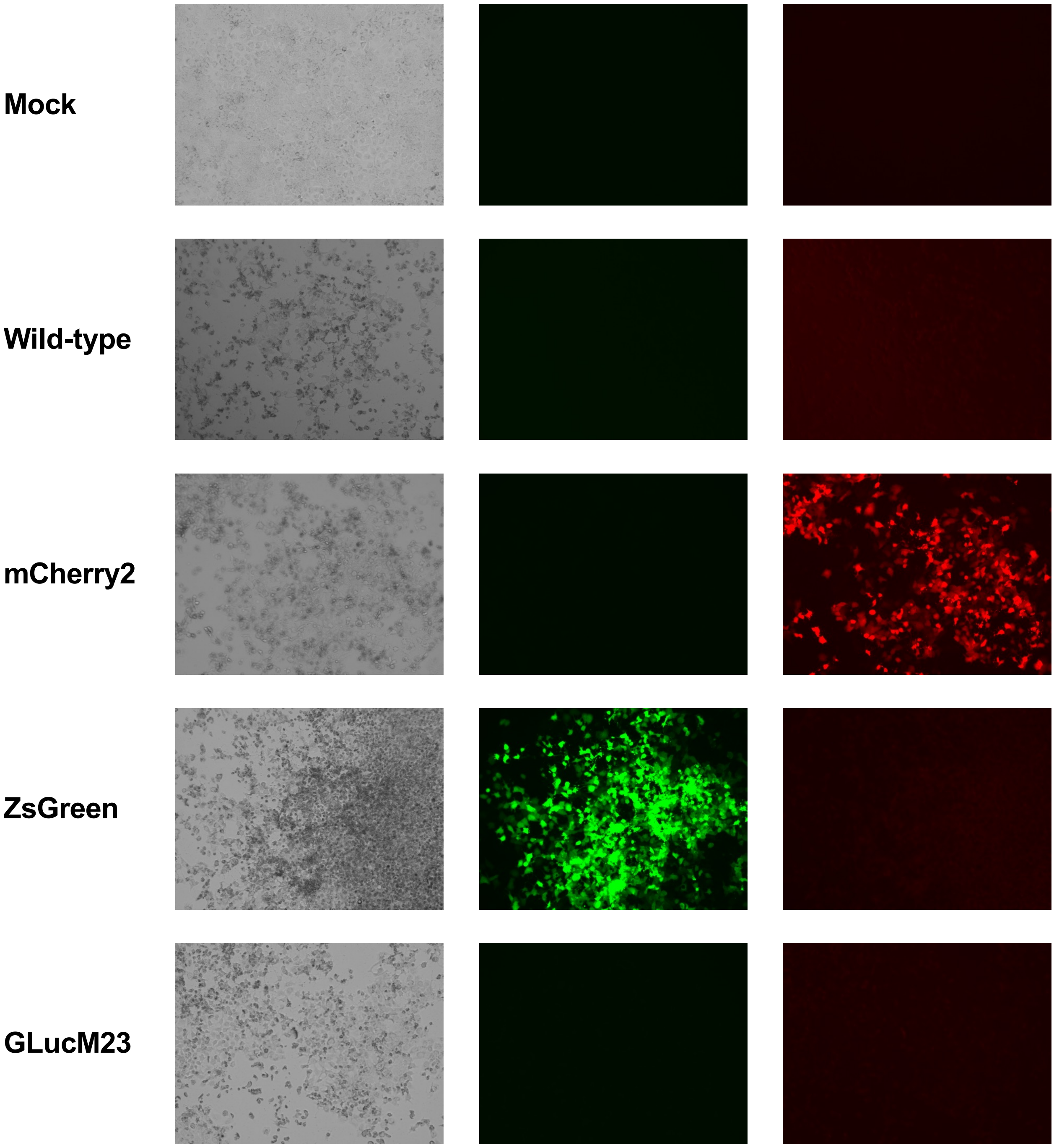

#### Supplementary Figure 3

**A** 15 hpi in NB-1 cells

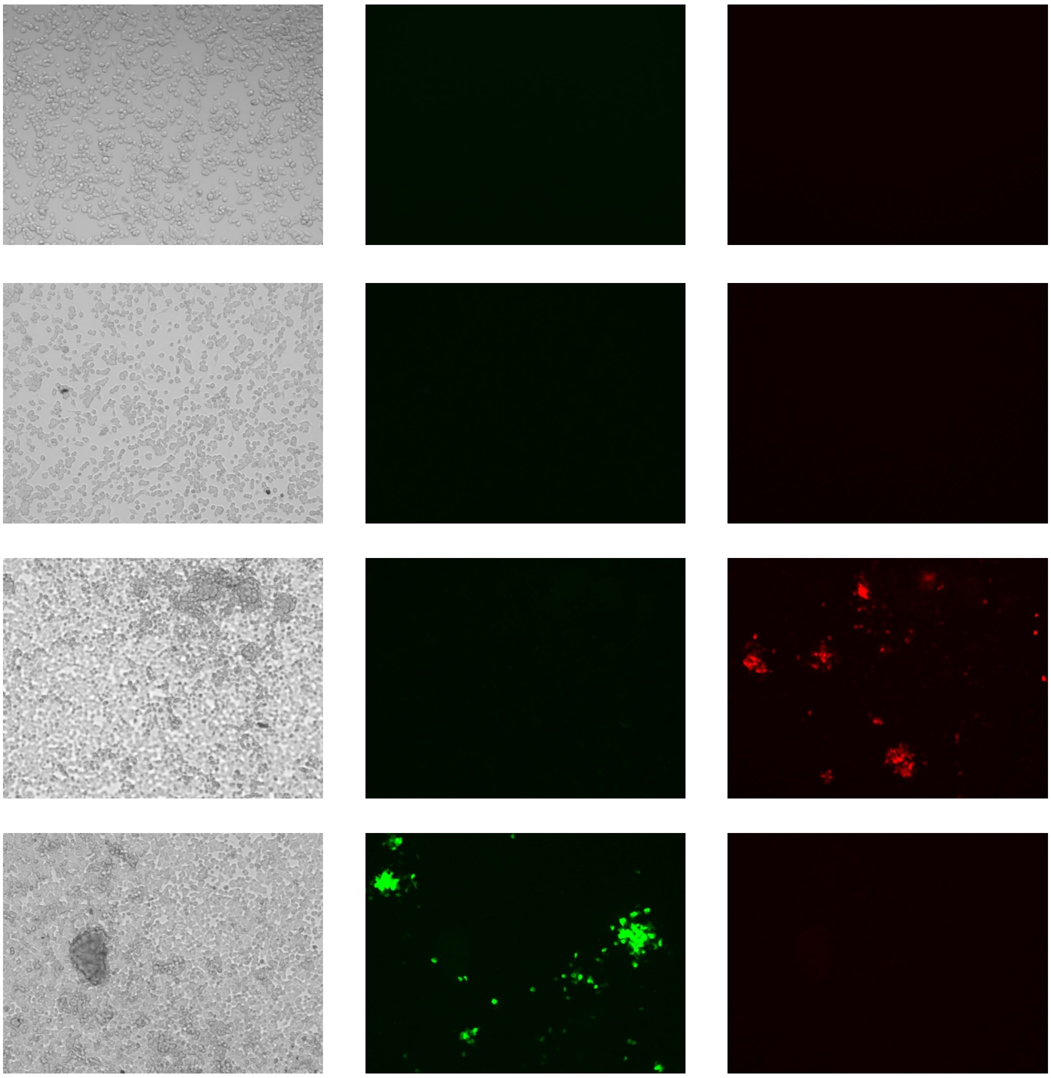

Mock

Wild-type

mCherry2

ZsGreen

#### Supplementary Figure 3

**B** 24 hpi in NB-1 cells

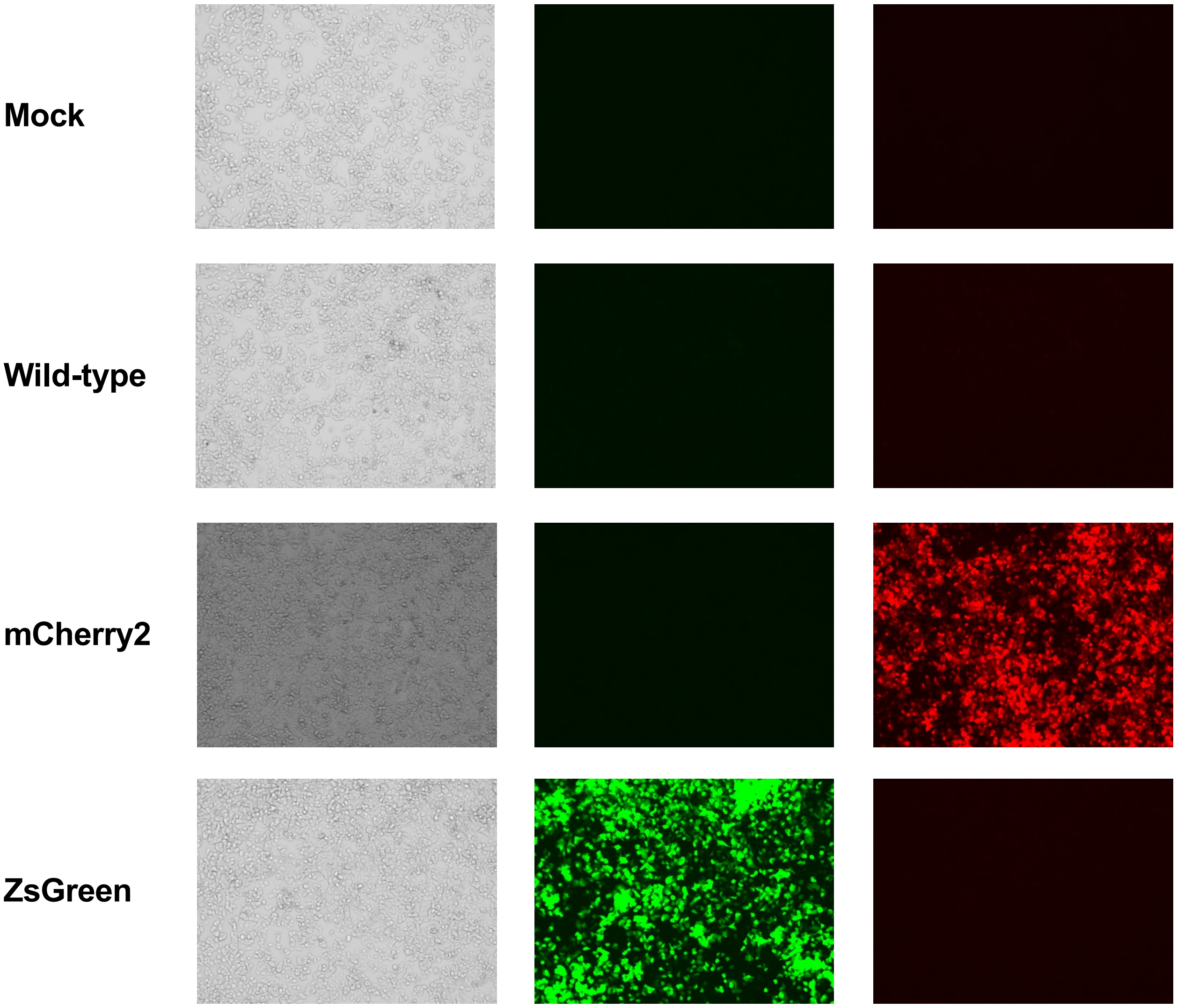

Supplementary Figure 4

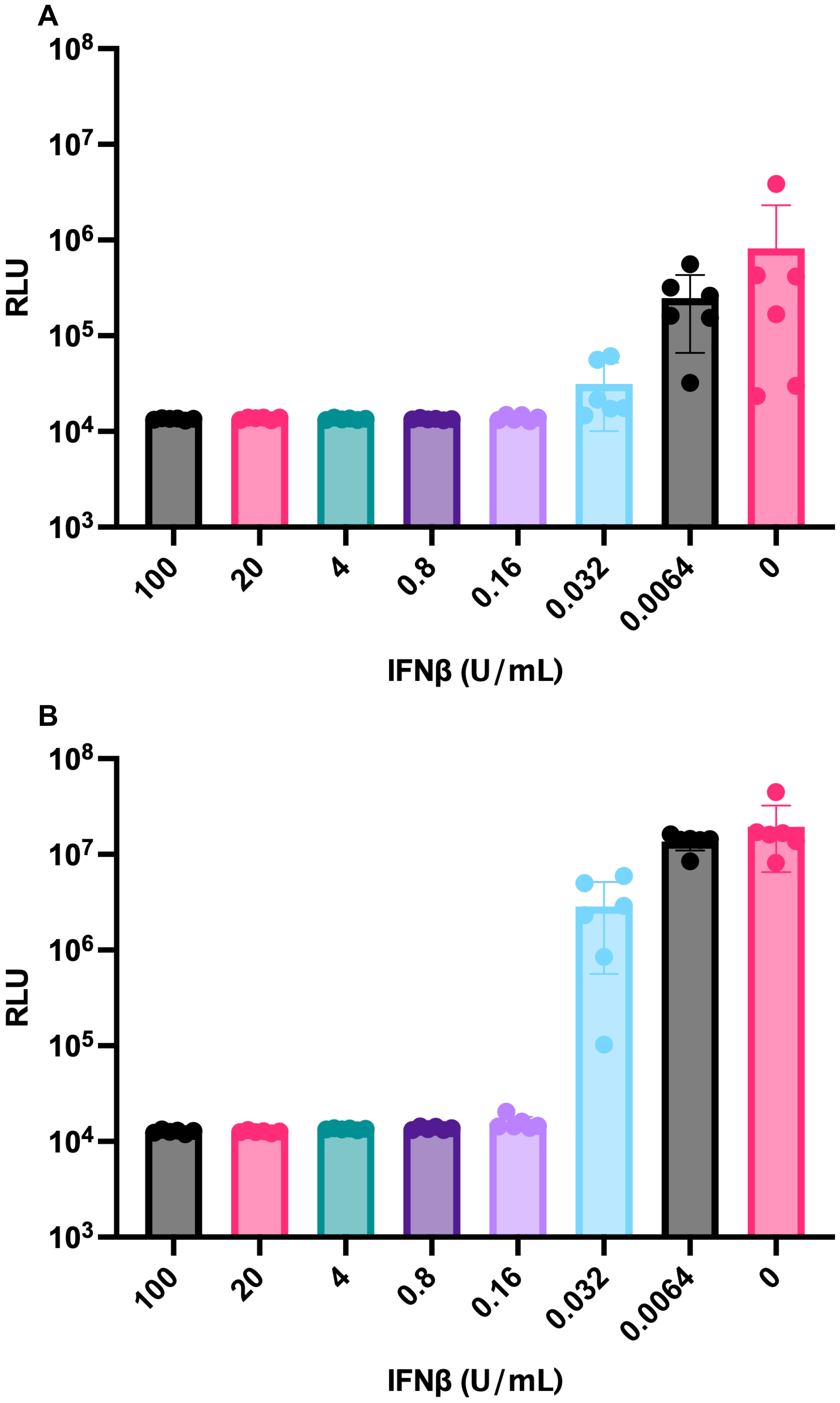

### Supplementary Figure 5

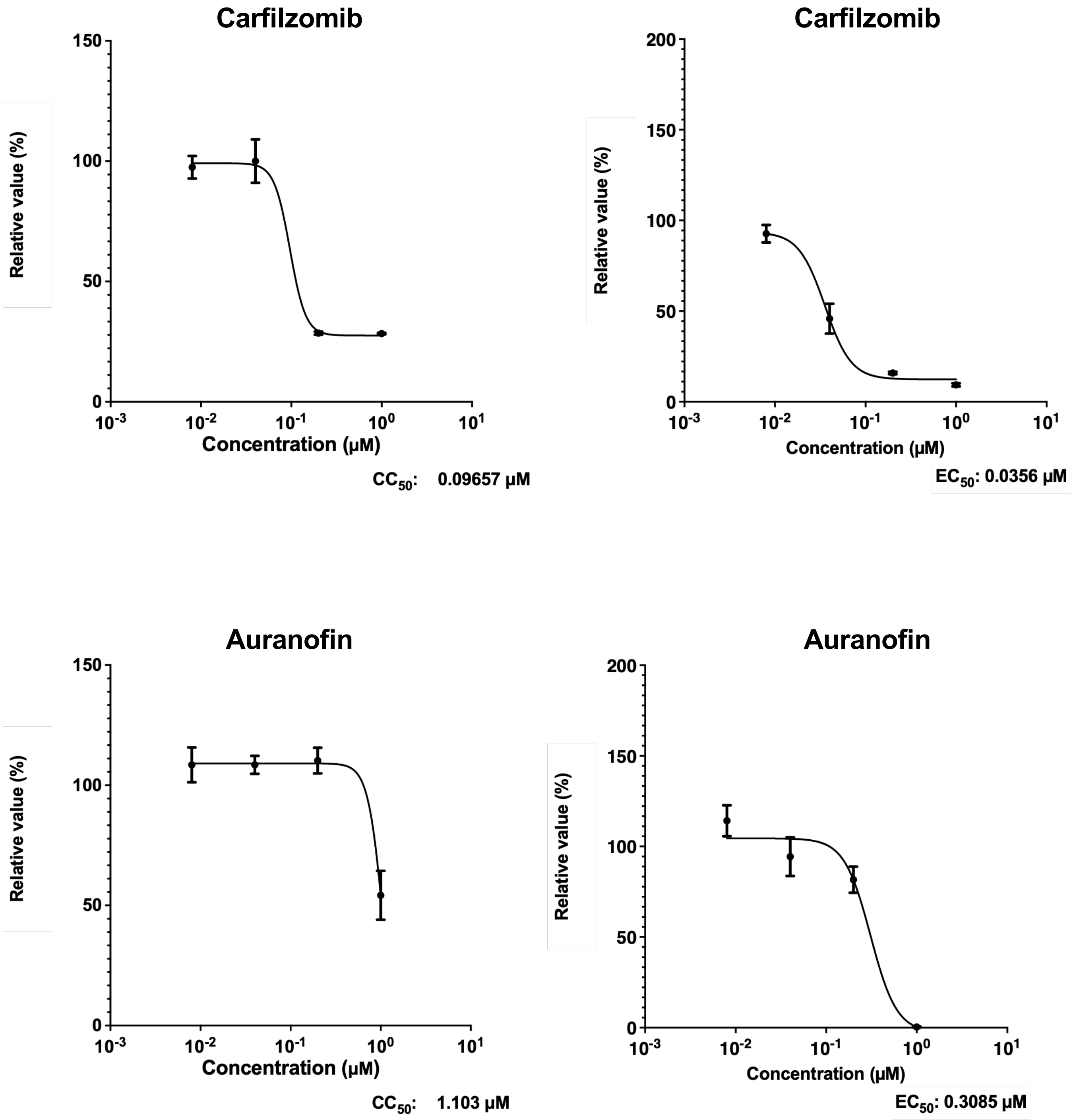

**Supplementary Table 1. Synthesized DNA sequences of the VEEV 3880 strain.**

The nucleotide sequences synthesized in this study are listed below. Red text indicates overlapping regions with the subsequent fragment.

| Fragment | Sequence |
|---|---|
| #1 | ATGGGCGGCGCATGAGAGAAGCCCAGACCAATTACCTACCCAAAATGGAGA<br>AAGTTCACGTTGACATCGAGGAAGACAGCCCATTCCTCAGAGCTTTGCAACG<br>GAGCTTCCCGCAGTTTGAGGTAGAAGCCAAGCAGGTCACTGATAATGACCAT<br>GCTAACGCCAGAGCGTTCTCGCATCTGGCTTCAAAACTGATCGAAACGGAGG<br>TGGACCCATCCGACACGATCCTTGACATTGGAAGTGCGCCCGCCCGCAGAAT<br>GTATTCTAAGCATAAGTACCATTGTATCTGTCCGATGAGATGTGCAGAAGAT<br>CCGGACAGATTGTACAAGTACGCAACTAAGCTGAAGAAAAACTGCAAGGAA<br>ATAACTGACAAGGAACTGGACAAGAAAATGAAAGAGCTCGCCGCCGTCATG<br>AGCGACCCTGACCTGGAAACTGAAACCATGTGCCTCCACGACGACGAGTCG<br>TGTCGCTACGAAGGGCAAGTCGCTGTTTACCAGGATGTATACGCGGTTGACG<br>GACCGACAAGTCTCTACCATCAAGCCAACAAGGGAGTTAGAGTCGCCTACT<br>GGATAGGCTTTGACACCACCCCTTTTATGTTCAAGAACTTGGCTGGAGCATA<br>TCCATCATACTCTACCAATTGGGCCGACGAAACCGTGTTAACGGCTCGTAAC<br>ATAGGCCTGTGCAGCTCCGACGTTATGGAGCGGTCACGTAGAGGGATGTCCA<br>TTCTTAGAAAGAAGTATCTGAAACCATCCAACAATGTTCTATTCTCTGTTGGC<br>TCGACCATCTACCACGAGAAGAGGGACTTACTGAGGAGCTGGCACCTGCCG<br>TCTGTATTTCACTTACGTGGCAAGCAAAATTACACATGTCGGTGTGAGACTA<br>TAGTTAGTTGCGACGGGTACGTCGTTAAAAGAATAGCTATCAGTCCAGGCCT<br>GTATGGGAAGCCTTCAGGCTATGCTGCTACGATGCACCGCGAGGGATTCTTG<br>TGCTGCAAAGTGACAGACACATTGAACGGGGAGAGGGTCTCTTTTCCCGTGT<br>GCACGTATGTGCCAGCTACACTGTGTGACCAAATGACTGGCATACTGGCAAC<br>AGATGTCAGTGCAGACGACGCACAAAAACTGCTGGTTGGGCTCAACCAGCG<br>TATAGTCGTCAACGGTCGCACCCAAAGAAACACCAATACCATGAAGAATTA<br>CCTTTTGCCAGTAGTAGCCCAGGCATTTGCTAGGTGGGCAAAGGAATATAAG<br>GAAGATCAAGAAGATGAGAGGCCGCTAGGACTACGAGACAGACAGTTAGTC<br>ATGGGGTGCTGTTGGGCTTTTAGAAGACACAAGATAACATCTATTTATAAAC<br>GCCCGGATACCCAAACCATCATCAAAGTGAACAGCGATTTCCACTCATTCGT<br>GCTGCCTAGAATAGGCAGTAACACATTGGAGATCGGGCTGAGGACAAGAAT<br>CAGGAAAATGTTAGAAGAGCACAAGGAGCCGTCACCTCTCATTACTGCCGA<br>GGACATACAAGAAGCTAAGTGCGCAGCCGACGAGGCTAAGGAGGTACGTGA<br>AGCCGAGGAGTTGCGTGCAGTACTACCACCTTTGGCAGCTGATGTTGAGGAG<br>CCCACTTTGGAAGCCGATGTCGACTTGATGTTACAAGAGGCTGGGGCCGGCT<br>CAGTGGAGACACCGCGTGGCTTGATAAAGGTTACCAGCTACGCCGGCGAGG<br>ACAAGATAGGCTCCTACGCAGTGCTTTCTCCGCAGGCCGTACTCAAGAGTGA |
| #2 | CTCCGCAGGCCGTACTCAAGAGTGAAAAACTATCTTGTATCCACCCCCTCGC<br>TGAACAAGTCATAGTGATAACACACTCTGGCCGAAAAGGGCGTTATGCCGT<br>GGAACCCTATCATGGAAAAGTAGTTGTGCCAGAGGGACACGCAATACCCGT<br>CCAGGATTTTCAAGCTCTGAGCGAAAGTGCTACCATTGTGTATAACGAGCGA |

| | |
|---|---|
| | GAGTTCGTAAACAGGTACCTGCACCATATTGCCACACACGGAGGAGCGCTG<br>AACACAGATGAAGAATACTATAAAACTGTCAAGCCCAGCGAGCACGACGGC<br>GAATACCTGTATGACATCGACAGGAAACAGTGCGTCAAGAAAGAACTAGTC<br>ACTGGGCTAGGGCTCACAGGCGAACTGGTGGATCCACCCTTCCATGAATTTG<br>CCTATGAGAGTCTGAGAACACGACCTGCCGCTCCCTACCAAGTACCAACCAT<br>AGGGGTGTATGGCGTGCCAGGATCAGGCAAGTCTGGCATCATCAAAAGCGC<br>GGTCACCAAAAAGGACCTGGTGGTGAGCGCCAAGAAAGAAAACTGTGCAGA<br>AATAATAAGGGACGTCAAGAGAATAAAAGGGCTGGACGTCAATGCCAGAAC<br>TGTGGACTCGGTGCTCCTGAATGGATGCAAATACCCCGTAGAGACCTTGTAT<br>ATTGACGAAGCTTTTGCTTGTCATGCAGGTACTCTTAGAGCGCTCATAGCCAT<br>CATAAGACCTAAAAAGGCAGTGCTCTGCGGGGATCCCAAACAGTGCGGCTT<br>TTTCAACATGATGTGCCTGAAAGTGCATTTTAATCATGAGATTTGCACGCAA<br>GTCTTCCATAAAAGCATCTCTCGCCGTTGCACTAAATCTGTGACTTCGGTCGT<br>CTCAACTTTGTTTTACGACAAAAGGATGAGAACGACAAACCCGAAAGAGAC<br>TAAGATTGAGATTGACACTACCGGCAGTACTAAACCCAAGCAGGACGATCT<br>CATCCTCACTTGTTTCAGAGGGTGGGTGAAGCAGTTGCAAATAGACTACAAA<br>GGCAACGAAGTAATGACGGCAGCTGCCTCCCAAGGGCTGACCCGTAAAGGA<br>GTGTATGCCGTTCGGTACAAGGTGAATGAAAATCCTCTTTACGCACCCACCT<br>CAGAACATGTGAACGTCCTACTGACCCGCACGGAGGACCGCATCGTGTGGA<br>AAACATTAGCCGGTGACCCATGGATAAAAACACTGACGGCCAAGTATCCTG<br>GGAATTTCACTGCCACGATAGAAGAATGGCAAGCAGAGCATGATGCCATCA<br>TGAGGCACATCTTGGAGAGACCGGACCCCACCGACGTCTTCCAAAACAAGG<br>CGAACGTGTGCTGGGCCAAGGCTCTAGTGCCGGTACTGAAGACTGCAGGTAT<br>AGACATGACCACCGAACAATGGAACACTGTGGATTACTTTGAAACGGACAA<br>AGCTCACTCAGCAGAGATAGTATTGAACCAATTATGCGTGAGGTTCTTTGGA<br>CTTGACCTGGACTCCGGTCTTTTCTCTGCACCCACAGTTCCGTTATCCATTAG<br>GAATAATCACTGGGATAACTCCCCGTCGCCTAACATGTACGGGCTGAATAAG<br>GAAGTGGTCCGTCAGCTCTCCCGCAGGTACCCACAACTGCCTCGTGCAGTTA<br>CCACTGGAAGAGTTTATGACATGAATACTGGTACACTGCGCAATTATGATCC<br>GCGCATAAATTTAGTACCTGTGAACAGAAGACTGCCTCACGCTTTAGTCCTC<br>CACCATAAT |
| #3 | TCACGCTTTAGTCCTCCACCATAATGAACACCCACAGAGTGACTTTTCTTCAT<br>TCGTCAGCAAATTGAAGGGCAGAACTGTCCTGGTGGTCGGGGAGAAGCTGT<br>CCGTCCCAGGCAAAACGGTAGACTGGCTGTCAGATCGGCCTGAGGCTACCTT<br>CAGAGCTCGGCTGGATTTAGGCATCCCGGGTGACGTGCCCAAATATGACATA<br>ATATTTATTAATGTAAGGACCCCATATAAATACCATCATTACCAGCAGTGCG<br>AAGACCATGCCATTAAGCTTAGCATGTTGACCAAGAAAGCTTGTCTGCATCT<br>GAATCCCGGCGGAACCTGTGTCAGCATAGGTTATGGTTACGCCGACAGGGCC<br>AGCGAAAGCATCATCGGCGCTATAGCGCGGCAGTTCAAGTTCTCCCGGGTAT<br>GCAAGCCGAAATCTTCACTTGAAGAGACAGAAGTTCTGTTTGTATTCATTGG<br>GTACGATCGCAAGGCCCGTACGCACAATCCTTACAAGCTTTCATCTACCCTG<br>ACCAACATCTATACAGGTTCCAGACTCCACGAAGCCGGATGCGCACCCTCAT<br>ATCATGTGGTGCGAGGGGACATTGCCACGGCCACCGAAGGAGTGATAATAA<br>ATGCTGCTAACAGCAAGGGACAACCTGGCGGAGGGGTGTGCGGAGCGCTGT<br>ACAAAAAATTCCCGGAAAGCTTCGATTTACAGCCGATCGAAGTGGGAAAAG<br>CGCGACTGGTCAAAGGCGCAGCTAAACATATTATTCATGCTGTAGGGCCGAA |

| | |
|---|---|
| | CTTCAATAAAGTCTCGGAAATTGAAGGTGACAAACAGTTGGCAGAGGCTTAT<br>GAGTCTATCGCTAAAATTGTCAACGACAACAATTACAAGTCAGTAGCGATTC<br>CACTGTTGTCCACCGGCATCTTTTCCGGGAACAAAGACCGCCTAACTCAATC<br>ATTGAACCATTTGCTGACAGCTTTAGACACCACCGATGCAGATGTAGCCATA<br>TACTGCAGGGACAAGAAATGGGAAATGACTCTCAAGGAAGCAGTGGCTAGG<br>AGAGAAGCAGTGGAGGAGATATGCATATCTGACGATTCATCGGTGACAGAA<br>CCGGATGCGGAGCTGGTGAGGGTGCATCCGAAGAGTTCTTTGGCAGGAAGG<br>AAAGGCTACAGCACAAGTGATGGCAAGACTTTCTCGTATTTGGAAGGGACTA<br>AATTCCACCAGGCGGCCAAGGATATAGCGGAAATTAATGCCATGTGGCCCG<br>TTGCAACGGAGGCCAATGAGCAAGTATGTATGTATATCCTCGGAGAAAGTAT<br>GAGCAGCATCAGGTCGAAATGCCCCGTCGAAGAGTCGGAAGCATCCACACC<br>GCCTAGCACGCTGCCTTGCTTGTGCATCCATGCTATGACTCCAGAAAGAGTA<br>CAGCGCCTGAAAGCGTCACGCCCAGAACAAATTACTGTGTGCTCATCCTTTC<br>CATTGCCGAAGTACAGAATCACTGGTGTGCAGAAGATCCAGTGCTCCCAGCC<br>TATACTGTTCTCACCGAAAGTGCCTGCATACATCCATCCACGGAAGTATCTC<br>GTGGAAACACCAACGGTAGAAGAGAACCAATCCACAGAGGGGACACCTGA<br>ACAACCGACACTTATAACCGTGGGTGAAACCAGAACTAGAACGCCCGAACC<br>GATTATTATTGAAGAAGAAGAGGATAGCATAAGCTTGCTGTCAGACGGCCC<br>GACCCACCAGGTGCTGCAAGTCGAGGCAGACATTCATGGGCCGCCTTCCGCA<br>TCCA |
| #4 | ATTCATGGGCCGCCTTCCGCATCCAGCTCATCTTGGTCCATTCCTCATGCATC<br>CGACTTTGATGTGGATAGTCTGTCCATCCTTGACACCCTGGAGGGAGCTAGC<br>GTGACCAGCGAAGAGGCATCAGTAGAGACTAACTCTCACTTCGCAAGGAGT<br>ATGGAGTTTCTGGCGCGACCGGTGCCTGCGCCCCGGACAGTATTCAGGAACC<br>CTCCACAGCCAGCTCCGCGCACAAGAACGCCGTCACTCGCACCTAGCAGGG<br>CCAGCTCGAGAATTAGCCTAGTTTCCAACCCGCCAGGAGTGAATAGGGTGAT<br>TACTAGGGAGGAGCTCGAGGCGCTCACCCCGTCACGTGCTCCCAGCAGGTCG<br>GTCTCAAGAACCAGCCTGGTTTCCAATCCTCCAGGCGTAAATAGGGTGATTA<br>CAAGAGAGGAGTTCGAGGCGTTCGTAGCACAACAACAATGACGGTTTGATG<br>CGGGTGCATACATCTTTTCCTCCGATACCGGTCAAGGGCATTTACAACAAAA<br>ATCAGTAAGGCAAACGGTGCTATCTGAAGTGGTGTTGGAGAGGACCGAATT<br>GGAGATTTCGTATGCCCCGCGCCTCGACCAAGAAAAAGAAGAGTTATTACG<br>CAAGAAATTACAGTTGAACCCCACACCCGCTAACAGAAGCAGATACCAGTC<br>CAGGAGAGTGGAGAACATGAAGGCCATAACAGCTAGACGTATTCTGCAAGG<br>CCTAGGGCATTATTTGAAGGCAGAAGGAAAAGTGGAGTGCTATCGAACCCT<br>GCATCCTGTTCCTTTGTATTCGTCCAGCGTGAATCGTGCCTTTTCAAGCCCCA<br>AGGTCGCAGTGGAAGCCTGCAACGCTATGTTGAAAGAAAACTTTCCGACCGT<br>GGCTTCTTACTGCATTATTCCGGAGTACGATGCCTATCTGGACATGGTTGACG<br>GCGCTTCTTGTTGTTTAGACACTGCCAGTTTTTGCCCTGCAAAGCTGCGCAGC<br>TTTCCAAAGAAACATTCCTATTTGGAACCCACAATACGATCGGCAGTGCCAT<br>CAGCGATTCAGAACACGCTCCAGAACGTCCTGGCAGCTGCCACGAAAAGAA<br>ATTGCAACGTCACGCAAATGAGAGAATTGCCCGTATTGGATTCGGCCGCCTT<br>TAATGTGGAATGCTTCAAGAAGTATGCATGCAATAATGAATATTGGGAAACG<br>TTTAAAGAAAACCCCATCAGGCTTACTGAAGAAAATGTGGTAAATTACATTA<br>CTAAATTAAAAGGACCAAAAGCTGCAGCTCTTTTTGCGAAGACACATAACTT<br>GAATATGTTACAGGACATACCAATGGATAGGTTTGTAATGGACTTGAAGAGG |

| | |
|---|---|
| | GATGTGAAAGTGACTCCAGGAACAAAACATACTGAGGAACGACCAAAGGTA<br>CAGGTGATCCAGGCTGCCGATCCGCTAGCAACAGCGTATCTGTGCGGAATAC<br>ACCGAGAGCTGGTCAGGAGATTAAACGCTGTCCTGCTTCCAAACATCCATAC<br>ACTGTTCGACATGTCAGCTGAAGACTTTGACGCTATTATCGCCGAGCATTTC<br>CAGCCTGGGGACTGCGTTCTGGAAACTGACATCGCGTCGTTTGACAAAAGCG<br>AGGATGACGCCATGGCTCTGACCGCGTTAATGATTCTGGAAGACTTAGGTGT<br>GGACGCAGAGCTGTTGACGCTGATCGAGGCGGCTTTCGGCGAGATTTCATCA<br>ATACATTTGCCCACTAAAACTAAATTTAAATTTGGAGCCATGATGAAATCTG |
| #5 | AAATTTGGAGCCATGATGAAATCTGGGATGTTCCTCACATTGTTTGTGAACA<br>CAGTCATCAACATCGTGATCGCAAGCAGAGTGTTAAGAGAACGGCTAACCG<br>GATCACCGTGTGCAGCATTTATTGGAGATGACAATATCGTGAAAGGAGTCAA<br>ATCGGACAAATTAATGGCAGACAGGTGCGCCACCTGGTTGAACATGGAAGT<br>CAAGATCATAGATGCTGTAGTGGGCGAGAAGGCGCCCTATTTCTGTGGAGGG<br>TTCATTTTGTGTGACTCCGTGACCGGCACAGCATGCCGTGTGGCAGACCCCT<br>TAAAAAGGCTGTTTAAGCTTGGCAAACCACTGGCAGCAGACGATGAACATG<br>ATGATGACAGGAGAAGGGCATTACATGAAGAGTCAACACGCTGGAACCGAG<br>TGGGAATTCTTCCAGAGCTGTGTAAGGCTGTAGAATCAAGGTATGAAACCGT<br>AGGAACTTCCATCATAGTTATGGCCATGACTACCTTAGCTAGCAGTGTTAAA<br>TCATTCAGCTACCTAAGAGGGGCCCCTATAACTCTCTACGGCTAACCTGAAT<br>GGACTACGACATAGTCTAGTCCGCCAAGATGTTCCCGTTCCAACCAATGTAT<br>CCTATGCAGCCAATGCCCTACCGCAACCCGTTCGCGGCCCCGCGCAGGCCCT<br>GGTTCCCCAGAACCGACCCTTTTCTGGCGATGCAGGTGCAGGAACTAACCCG<br>CTCGATGGCCAACCTGACGTTCAAGCAACGCCGGGAAGCGCCGCCTGAGGG<br>GCCACCTGCTAAGAAACCGAAGAGGGAGGCCCCGCAAAAACAAAAAGGGG<br>GGGGCCAAGGGAAGAAGAAGAAGAACCAGGGGAAGAAGAAGGCTAAGACG<br>GGACCGCCCAATCCGAAGGCACAAAATGGAAACAAGAAGAAGACCAACAA<br>GAAACCAGGCAAGAGACAACGCATGGTTATGAAACTGGAATCTGATAAGAC<br>ATTCCCAATTATGCTGGAAGGAAAGATTAACGGCTACGCTTGCGTGGTCGGA<br>GGGAAGTTATTCAGGCCGATGCATGTGGAAGGCAAAATCGATAACGATGTT<br>CTGGCCGCGCTTAAGACGAAGAAAGCGTCCAAATATGATCTTGAGTATGCA<br>GATGTGCCACAGAACATGCGGGCCGATACATTCAAGTACACCCATGAGAAA<br>CCCCAAGGCTACTACAGCTGGCACCACGGAGCAGTCCAATATGAAAACGGG<br>CGTTTCACGGTGCCAAAAGGAGTTGGAGCTAAAGGAGACAGTGGGCGACCC<br>ATTCTGGATAATCAGGGACGGGTGGTCGCTATTGTGCTAGGAGGTGTGAACG<br>AAGGATCTAGGACAGCCCTTTCAGTCGTCATGTGGAACGAGAAGGGAGTTA<br>CTGTGAAGTATACTCCGGAGAATTGCGAGCAATGGTCACTAGTGACCACCAT<br>GTGCCTGCTCGCCAATGTGACGTTCCCATGCGCTCAACCACCAATTTGCTAC<br>GACAGAAAACCAGCAGAGACCTTGGCCATGCTCAGCGTCAACGTGGACAAC<br>CCGGGCTACGACGAGCTGCTAGAAGCAGCTGTTAAGTGCCCAGGAAGGAAA<br>AGGAGATCTACTGAGGAGCTGTTTAAGGAGTATAAGCTAACGCGTCCCTACA<br>TGGCCAGATGTATTAGGTGCGCCGTTGGGAGCTGCCATAGTCCAATAGCAAT<br>TGAGGCAGTGAAGAGCGACGGGCACGACGGCTATGTCAGACTCCAGACTTC<br>CTCACA |
| #6 | ATGTCAGACTCCAGACTTCCTCACAGTATGGCCTGGATTCTTCCGGCAACTT<br>AAAGGGCAGGACCATGCGGTATGACATGCACGGGACCATTGAAGAGATACC<br>ACTACATCAAGTGTCACTTCATACATCTCGCCCGTGTCACATTGTGGATGGG |

| | |
|---|---|
| | CATGGCTACTTTTTGCTTGCTAGATGCCCGGCAGGGGATTCCATCACCATGG<br>AATTTAAGAAAGATGCAGTCACACACTCCTGCTCAGTGCCGTATGAAGTGAA<br>ATTTAATCCTGTAGGCAGAGAACTCTACACTCACCCCCCAGAACACGGAGCA<br>GAGCAAGCGTGCCAAGTCTACGCACATGATGCACAGAACAGAGGAGCTTAT<br>GTCGAGATGCACCTCCCGGGCTCTGAAGTGGACAGCAGTCTGGTCTCTTTGA<br>GCGGCAGTTCAGTCACTGTGACACCTCCTGCGGGGACTAGCGCCCTGGTGGA<br>ATGCGAGTGCGGTGGCACAAAGATCTCCGAGACCATCAACACGGCAAAACA<br>GTTCAGCCAGTGCACAAAGAAGGAGCAGTGCAGAGCATATCGACTGCAGAA<br>TGACAAGTGGGTGTATAATTCTGACAAACTGCCCAAAGCAGCGGGAGCCAC<br>TCTAAAAGGAAAACTACATGTCCCATTCTTGCTGGCAGACGGCAAATGCACC<br>GTGCCTCTAGCACCGGAACCGATGATAACCTTCGGATTCCGATCAGTGTCAC<br>TGAAACTGCACCCCAAGAATCCCACATATCTGACTACCCGCCAACTTGCTGA<br>TGAGCCTCATTACACGCACGAGCTCATATCTGAACCGGTTGTTAGGAACTTT<br>TCTGTCACTGAAAAAGGATGGGAGTTTGTATGGGGTAACCACCCGCCGAAA<br>AGGTTTTGGGCACAGGAAACAGCACCCGGGAATCCACATGGGCTGCCGCAC<br>GAGGTGATAACTCATTATTACCACAGATACCCCATGTCCACTATCCTGGGTT<br>TGTCAATTTGCGCCGCCATCGTGACCGTTTCCATCGCAGCGTCCACCTGGCTG<br>TTATGCAAATCCAGAGTCTCGTGCCTAACTCCTTACCGGCTAACACCTAACG<br>CCAGGATGCCGCTTTGCCTGGCTGTGCTTTGCTGTGCTCGCACTGCCCGGGCC<br>GAGACCACCTGGGAGTCCTTGGACCATCTATGGAATAATAACCAACAGATGT<br>TCTGGATCCAATTGCTGATCCCTCTGGCCGCCTTGATTGTAGTGACTCGCCTG<br>CTCAGGTGTGTGTGCTGTGTCGTGCCTTTTTTAGTCGTGGCCGGCGCCGCAGG<br>CGCCGGCGCCTACGAGCACGCGACCACGATGCCGAGCCAAGCGGGAATCCC<br>GTATAACACCATAGTCAACAGAGCAGGCTACGCACCACTCCCTATCAGCATA<br>ACACCAACAAAGATCAAGCTGATACCCACAGTGAACTTGGAGTACGTCACC<br>TGCCACTACAAGACAGGAATGGATTCACCAGCCATTAAATGCTGCGGATCGC<br>AGGAATGCACTCCAACTTACAGGCCTGATGAACAGTGCAAAGTCTTTACAGG<br>GGTTTACCCATTCATGTGGGGAGGGGCATATTGCTTTTGCGACACTGAGAAT<br>ACCCAAGTCAGCAAGGCTTATGTAATGAAATCTGACGACTGCCTTGCGGATC<br>ATGCTGAAGCATACAAAGCGCACACAGCCTCAGTGCAGGCGTTCCTCAACAT<br>CACAGTGGGAGAACATTCCATCGTGACCACCGTGTATGTGAATGGAGAAAC |
| #7 | CCACCGTGTATGTGAATGGAGAAACTCCTGTGAACTTCAATGGGGTCAAATT<br>AACTGCAGGCCCACTTTCCACAGCTTGGACACCCTTTGACCGCAAAATTGTG<br>CAGTATGCCGGGGAGATCTACAATTACGATTTTCCTGAGTACGGGGCAGGAC<br>AGCCAGGAGCTTTTGGAGACATACAATCCAGAACAGTTTCAAGCTCAGATCT<br>GTACGCCAACACCAACCTAGTGCTGCAGAGACCCAAAGCAGGAGCGATCCA<br>TGTGCCATACACACAGGCACCATCGGGTTTTGAGCAATGGAAGAAAGATAA<br>GGCTCCATCATTGAAATTTACTGCCCCTTTTGGATGTGAAATATATACAAACC<br>CCATTCGCGCCGAAAATTGTGCTGTGGGGTCAATTCCATTAGCCTTTGACAT<br>CCCAGACGCCTTGTTCACCAGGGTGTCAGAAACACCGACACTATCAGCGGCC<br>GAATGCACTCTTAATGAGTGCGTGTATTCATCCGACTTTGGCGGGATCGCCA<br>CAGTCAAGTATTCGGCCAGCAAGTCAGGCAAATGCGCAGTCCATGTGCCATC<br>AGGGACAGCCACCCTAAAAGAAGCAGCAATCGAGTTAGCCGAGCAAGGGTC<br>GGCGACTATTCATTTCTCGACTGCAAACATCCACCCGGAGTTCAGGCTCCAA<br>ATATGCACATCATACGTTACGTGCAAAGGTGATTGTCACCCCCCGAAAGACC<br>ATATTGTGACACACCCCCAGTATCACGCCCAAACATTTACAGCCGCGGTGTC |

| | |
|---|---|
| | AAAAACCGCGTGGACGTGGTTAACATCCCTGCTGGGAGGATCAGCTGTAATT<br>ATTATAATTGGCTTAGTGCTGGCTACTATTGTGGCCATGTACGTGCTGACCAA<br>CCAGAAACA<u>TAA</u>TTGAACACAGCAGCAATTGGCAAGCTGCTCATATAGAAT<br>TTGCGGCGATTGGAATGCCGCTTTTAAAATTTTTATTTTATTTTCTTTTCTTTT<br>CCGAATCGGATTTTGTTTTTAATATTTC |

**Supplementary Table 2. Primers used for sequencing the pCMV-VEEV 3880 plasmid.**

The primers used for full-length sequencing of the pCMV-VEEV 3880 plasmid are listed below.

| Name of primer | Nucleotide sequence |
|---|---|
| 3880-Seq1 | CCCTGACCTGGAAACTGAAA |
| 3880-Seq2 | GAGGGTCTCTTTTCCCGTGT |
| 3880-Seq3 | GAAGCCGATGTCGACTTGAT |
| 3880-Seq4 | CGCTCCCTACCAAGTACCAA |
| 3880-Seq5 | GTTTCAGAGGGTGGGTGAAG |
| 3880-Seq6 | GAAGTGGTCCGTCAGCTCTC |
| 3880-Seq7 | ATGCAAGCCGAAATCTTCAC |
| 3880-Seq8 | ACACCACCGATGCAGATGTA |
| 3880-Seq9 | CTGTGTGCTCATCCTTTCCA |
| 3880-Seq10 | AGGGCCAGCTCGAGAATTAG |
| 3880-Seq11 | ACCGTGGCTTCTTACTGCAT |
| 3880-Seq12 | CAGCGTATCTGTGCGGAATA |
| 3880-Seq13 | AGAAGGCGCCCTATTTCTGT |
| 3880-Seq14 | CAACCTGACGTTCAAGCAAC |
| 3880-Seq15 | ACAGCCCTTTCAGTCGTCAT |
| 3880-Seq16 | ATGGGCATGGCTACTTTTTG |
| 3880-Seq17 | AGCGGGAGCCACTCTAAAAG |
| 3880-Seq18 | TCGTGCCTAACTCCTTACCG |
| 3880-Seq19 | AGCATACAAAGCGCACACAG |

**Supplementary Table 3. Synthesized sequences for the construction of reporter viruses.**

The nucleotide sequences synthesized in this study are listed below. Colored text indicates the coding sequences for each reporter protein.

| **Fragment** | **Sequence** |
|---|---|
| mCherry2 | GAATTGCGAGCAATGGTCACTAGTGACCACCATGGTTTCAAAGGGTGAGGAGGACAA<br>TATGGCTATTATCAAAGAGTTTATGAGGTTCAAGGTTCACATGGAAGGCTCAGTGAACG<br>GCCACGAGTTCGAAATTGAAGGCGAGGGTGAGGGTCGACCTTATGAAGGCACGCAAA<br>CGGCCAAACTGAAAGTGACAAAAGGTGGACCGCTGCCCTTCGCTTGGGACATCCTGA<br>GTCCGCAATTCATGTACGGTTCCAAGGCTTACGTCAAACACCCGGCTGATATCCCTGAT<br>TATCTGAAGCTGTCATTTCCAGAGGGGTTTAATTGGGAACGAGTAATGAACTTCGAAGA<br>CGGTGGTGTGGTTACAGTAACCCAAGACAGCTCCTTGCAGGACGGAGAATTCATTTAC<br>AAAGTCAAATTGCGCGGGACGAATTTTCCTAGCGACGGTCCAGTGATGCAATGTCGAA<br>CGATGGGATGGGAAGCATCCACGGAGAGGATGTACCCTGAGGATGGCGCGCTGAAGG<br>GGGAAATTAAGCAGCGGCTGAAATTGAAGGATGGAGGACATTATGATGCTGAGGTCAA<br>AACCACATATAAGGCTAAGAAGCCAGTACAGTTGCCTGGTGCATACAATGTTGACATCA<br>AGCTTGATATACTTTCTCATAATGAGGATTATACAATAGTCGAGCAATACGAGCGAGCGG<br>AAGGCCGGCACTCTACTGGAGGAATGGATGAACTCTATAAAACCGGTGGAAGCGGAG<br>CTACTAACTTCAGCCTGCTGAAGCAGGCTGGCGACGTGGAGGAGAACCCTGGACCTT<br>CACTAGTGACCACCATGTGCCTGCTCGCCAATGT |
| ZsGreen | GAATTGCGAGCAATGGTCACTAGTGACCACCATGGCCCAGTCCAAGCACGGCCTGAC<br>CAAGGAGATGACCATGAAGTACCGCATGGAGGGCTGCGTGGACGGCCACAAGTTCGT<br>GATCACCGGCGAGGGCATCGGCTACCCCTTCAAGGGCAAGCAGGCCATCAACCTGTGC<br>GTGGTGGAGGGCGGCCCCTTGCCCTTCGCCGAGGACATCTTGTCCGCCGCCTTCATGT<br>ACGGCAACCGCGTGTTCACCGAGTACCCCCAGGACATCGTCGACTACTTCAAGAACTC<br>CTGCCCCGCCGGCTACACCTGGGACCGCTCCTTCCTGTTCGAGGACGGCGCCGTGTGC<br>ATCTGCAACGCCGACATCACCGTGAGCGTGGAGGAGAACTGCATGTACCACGAGTCCA<br>AGTTCTACGGCGTGAACTTCCCCGCCGACGGCCCCGTGATGAAGAAGATGACCGACAA<br>CTGGGAGCCCTCCTGCGAGAAGATCATCCCCGTGCCCAAGCAGGGCATCTTGAAGGGC<br>GACGTGAGCATGTACCTGCTGCTGAAGGACGGTGGCCGCTTGCGCTGCCAGTTCGACA<br>CCGTGTACAAGGCCAAGTCCGTGCCCCGCAAGATGCCCGACTGGCACTTCATCCAGCA<br>CAAGCTGACCCGCGAGGACCGCAGCGACGCCAAGAACCAGAAGTGGCACCTGACCG<br>AGCACGCCATCGCCTCCGGCTCCGCCTTGCCCACCGGTGGAAGCGGAGCTACTAACTT<br>CAGCCTGCTGAAGCAGGCTGGCGACGTGGAGGAGAACCCTGGACCTTCACTAGTGAC<br>CACCATGTGCCTGCTCGCCAATGT |
| GLucM23 | GAATTGCGAGCAATGGTCACTAGTGACCACCATGGGAGTCAAAGTTCTGTTTGCCCTG<br>ATCTGCATCGCTGTGGCCGAGGCCAAGCCCACCGAGAACAACGAAGACTTCAACATC<br>GTGGCCGTGGCCAGCAACTTCGCGACCACGGATCTCGATGCTGACCGCGGGAAGTTG<br>CCCGGCGAGAAGCTGCCGCTGGAGGTGCTCAAAGAGCTTGAAGCCAATGCCCGGAAA<br>GCTGGCTGCACCAGGGGCTGTCTGATCTGCCTGTCCCACATCAAGTGCACGCCCAAGA<br>TGAAGAAGTTCATCCCAGGACGCTGCCACACCTACGAAGGCGACAAAGAGTCCGCAC<br>AGGGCGGCATAGGCGAGGCGATCGATGACATTCCTGAGATTCCTGGGTTCAAGGACT<br>TGGAGCCCATTGAGCAGTTCATCGCACAGGTCGATCTGTGTGTGGACTGCACAACTGG<br>CTGCCTCAAAGGGCTTGCCAACGTGCAGTGTTCTGACCTGCTCAAGAAGTGGCTGCCG<br>CAACGCTGTGCGACCTTTGCCAGCAAGATCCAGGGCCAGGTGGACAAGATCAAGGGG<br>GCCGGTGATGACACCGGTGGAAGCGGAGCTACTAACTTCAGCCTGCTGAAGCAGGCT |

| | |
|---|---|
| | GGCGACGTGGAGGAGAACCCTGGACCTTCACTAGTGACCACCATGTGCCTGCTCGCCAATGT |
